# Dynamical models reveal anatomically reliable attractor landscapes embedded in resting state brain networks

**DOI:** 10.1101/2024.01.15.575745

**Authors:** Ruiqi Chen, Matthew Singh, Todd S. Braver, ShiNung Ching

## Abstract

Analyses of functional connectivity (FC) in resting-state brain networks (RSNs) have generated many insights into cognition. However, the mechanistic underpinnings of FC and RSNs are still not well-understood. It remains debated whether resting state activity is best characterized as noise-driven fluctuations around a single stable state, or instead, as a nonlinear dynamical system with nontrivial attractors embedded in the RSNs. Here, we provide evidence for the latter, by constructing whole-brain dynamical systems models from individual resting-state fMRI (rfMRI) recordings, using the Mesoscale Individualized NeuroDynamic (MINDy) platform. The MINDy models consist of hundreds of neural masses representing brain parcels, connected by fully trainable, individualized weights. We found that our models manifested a diverse taxonomy of nontrivial attractor landscapes including multiple equilibria and limit cycles. However, when projected into anatomical space, these attractors mapped onto a limited set of canonical RSNs, including the default mode network (DMN) and frontoparietal control network (FPN), which were reliable at the individual level. Further, by creating convex combinations of models, bifurcations were induced that recapitulated the full spectrum of dynamics found via fitting. These findings suggest that the resting brain traverses a diverse set of dynamics, which generates several distinct but anatomically overlapping attractor landscapes. Treating rfMRI as a unimodal stationary process (i.e., conventional FC) may miss critical attractor properties and structure within the resting brain. Instead, these may be better captured through neural dynamical modeling and analytic approaches. The results provide new insights into the generative mechanisms and intrinsic spatiotemporal organization of brain networks.

## 1 Introduction

Resting state fMRI (rfMRI) has become an important tool to probe the link between ongoing brain dynamics and cognition. The most common analytic approach utilized in rfMRI studies is to characterize brain-wide statistical associations (known as functional connectivity or FC) and relate them to cognitive and behavioral indices. However, the dynamical processes that generate the observed rfMRI fluctuations and statistics (e.g., FC) remain elusive. In particular, it is unknown whether resting state dynamics can be best described as a unimodal stationary process featuring statistical fluctuations around the mean, or instead, as a nonlinear dynamic system with nontrivial fluctuations associated with stable patterns different from the mean.

The prior FC literature has provided mixed support for both hypotheses. Traditionally, FC is considered stationary over the scanning session (Biswal et al., 1995). Correspondingly, the underlying dynamics are found to contain a stable equilibrium (point attractor) at the global mean, and the noisy fluctuations around this stable mean produce the observed FC pattern (Deco, Ponce-Alvarez, et al., 2013). However, recent years have witnessed the rapid development of an analysis technique called time-varying functional connectivity (tvFC), also known as dynamic FC (Lurie et al., 2020). The tvFC method identifies recurring short-time-windowed FC patterns that differ from the mean FC. These transient patterns are reliable within individuals and across populations (Abrol et al., 2017; Choe et al., 2017). More interestingly, transient FC patterns but not the mean FC were found to predict psychopathology (Reinen et al., 2018). These findings seem to suggest the existence of nontrivial, functionally salient fluctuation in resting state dynamics.

However, the nature and validity of tvFC characterization is itself still under debate. For example, it is known that head motion and physiological noise generate confounds in FC (Power et al., 2012), and even more so for tvFC, which relies on data from shorter duration timeseries (i.e., windowed epochs). More fundamentally, even if tvFC faithfully captures the temporal evolution of neural covariation patterns, it is still unclear whether tvFC states are merely snapshots of the noisy fluctuations around a stable mean, or if they can be associated with nontrivial dynamics. Indeed, tvFC states might be generated from various kinds of nontrivial dynamics (Heitmann & Breakspear, 2018). However, an influential paper (Laumann et al., 2017) showed that tvFC clustering would produce very similar results when applied to either real data or stationary noise with matched mean FC and power spectral density. Therefore, to understand the substrate of brain-wide associations, their temporal fluctuations, and ultimately the resting state dynamics that produce such associations, it is necessary to go beyond descriptive methods like tvFC clustering and adopt a more mechanistic framework.

Dynamical systems modeling and analysis can provide unique insights into the problem of nontrivial fluctuations in resting state dynamics. Dynamic models of brain activity predict the evolution of activation timeseries given an initial estimate of hidden states. They thus provide a generative mechanism for resting state dynamics and associated statistics, such as FC. To date, the evidence for nontrivial fluctuations from dynamic modeling is also mixed. There are two relevant types of neural models utilized to characterize resting state brain dynamics as measured by rfMRI: structural-connectome-informed models, and directly-parameterized models. The first type of models usually contain hundreds of sub-components representing brain parcels, connected by weights derived from the brain’s structural connectome, e.g., through diffusion tensor imaging. Early models typically included only a few free parameters, such as the global scaling factor for connectivity, that are directly fit to the fMRI data. It was consistently found that the emergent dynamics involved multiple nontrivial attractors (Deco & Jirsa, 2012; Deco, Ponce-Alvarez, et al., 2013), and the distribution of nontrivial attractors might reflect the organization of resting state functional networks (Golos et al., 2015). However, a recent study (Sip et al., 2023) that replaced the neural mass approximation of regional dynamics with a more powerful approximation scheme (i.e., an artificial neural network) reported the opposite, with a single globally stable attractor located at the mean. The second type of models do not assume that the structural connectome is a good surrogate for functional coupling, but rather directly optimize the effective connectivity between regions by predicting empirical fMRI time series. Most of the existing works adopting this approach has utilized the framework of Dynamical Causal Modeling (DCM) (Friston et al., 2003). Although nonlinear DCM has been proposed, it is computationally too expensive for more than ten nodes (Friston et al., 2019). Therefore, most rfMRI DCMs have used a linear approximation (Razi et al., 2017), which by definition cannot express nontrivial fluctuations. It has been argued that such stationary linear models have even lower mean estimation error than common nonlinear models for rfMRI (Nozari et al., 2023). However, a rigorous Bayesian model comparison found that a time-varying (‘dynamic’, short-time-windowed) linear DCM clearly outperformed a stationary linear DCM (Park et al., 2018). In short, previous studies have associated rfMRI with a variety of dynamics ranging from a monostable linear or weakly nonlinear system to a multistable strongly nonlinear system, with diverging evidence for nontrivial fluctuations. What might be the explanation for such inconsistent results?

Here, we suggest that prior approaches have captured some, but not all of the critical aspects of resting state brain dynamics. We hypothesize that the resting brain is particularly sensitive to modulation, and as such, can manifest a spectrum of different dynamics that systematically vary across individuals and time. It has been suggested that the resting brain is close to bifurcation, such that a small change in control parameters will alter the stability of the trivial attractor located at the mean (Deco, Ponce-Alvarez, et al., 2013). However, it remains unknown whether both sides of the bifurcation can be observed in a same fMRI dataset, and whether such a bifurcation best characterizes differences between individuals, or state changes within individuals across different time periods. Previous models were either too constrained to express diverse sets of dynamics, or lack the specificity to describe individual differences and session-to-session variations. In this project, we overcome these prior limitations by adopting the Mesoscale Individualized NeuroDynamics (MINDy) framework (Singh, Braver, et al., 2020). A key advantage of MINDy models is that they combine the expressiveness of nonlinear neural mass models with the flexibility and individuality of directly parameterized effective connectivity. Our prior work validated that MINDy models generate individualized, robust, and reliable fits of rfMRI data, with nontrivial dynamics observed (Singh, Braver, et al., 2020). Here, we used the MINDy framework to analyze the taxonomy of resting state brain dynamics, and to more comprehensively characterize how they change across individuals and time. We fit MINDy models of rfMRI data from over five hundred participants and each of two scanning sessions in the Human Connectome Project (HCP) (Smith et al., 2013) to elucidate the dynamic profiles that best explained rfMRI signals. We then analyzed the existence of anatomically reliable attractors and ghost attractors, which are the signatures of a class of bifurcations, showing that the latter frequently occurs. Finally, we show that such attractors were consistent across the population and represent differential activation of well-known functional brain networks, such as the default mode network (DMN) and frontoparietal control network (FPN).

## 2 Methods

### 2.1 Data preprocessing

We used the rfMRI data from the HCP Young Adult dataset (Smith et al., 2013). Informed consent was obtained during the original study. Researchers were required to sign the data use terms before accesssing the data. Data was originally collected on a 3T scanner with a TR of 720ms and 2mm isotropic voxels. Participants underwent two scanning sessions on separate days. Each session included two scanning runs of 1200 TRs (around 15 minutes), one using right-to-left phase encoding direction and the other left-to-right. Participants were instructed to stay awake with eyes open and relax fixation on a bright cross hair on a dark background, presented in a darkened room.

We adopted the preprocessing pipeline suggested in (Siegel et al., 2017), which was shown to effectively suppress the influence of head motion in rsFC-behavior associations. Because we are particularly interested in session-to-session variations in dynamics, we used a relatively strict inclusion criteria to make sure the data in all runs were sufficiently clean. Specifically, we only included participants with no missing runs or runs that had more than 1/3 (400 out of 1200) frames with high head motion (see below), resulting in a total number of 510 participants.

We began with the rfMRI data provided by HCP that has been minimally preprocessed, motion-corrected, and denoised with FIX-ICA (Smith et al., 2013). Following (Siegel et al., 2017), data was first detrended and then motion scrubbed with framewise displacement (FD) and temporal derivative of variation (DVARS) (Afyouni & Nichols, 2018). FD and DVARS were filtered for respiratory artifact with a 40-th order 0.06-0.14Hz band stop filter. Frames with FD above 0.2mm or DVARS above 1.05 times of the median were linearly interpolated. We then regressed out from the data the top five principal components of the white matter and the cerebrospinal fluid signals (CompCor), and the mean signal from the gray matter.

After preprocessing, the data was averaged within each parcel according to the 200-parcel atlas from (Schaefer et al., 2018). Data points exceeding 5 standard deviations in each time series were linearly interpolated. To obtain the underlying neural activity, we deconvolved the data with the canonical haemodynamic response function (HRF) from (Friston et al., 1998) using Wiener deconvolution (Wiener, 1949), a deconvolution technique that minimizes the influence of noise. We used a 30-point HRF kernel and a noise-power to signal-power ratio of 0.02. Finally, the data were z-scored within each timeseries.

### 2.2 Model architecture and fitting

We adopted the Mesoscale Individualized NeuroDynamics (MINDy) framework from (Singh, Braver, et al., 2020). A MINDy model contains interconnected neural masses representing brain parcels, with trainable and individualized connection weights. Each neural mass is assumed to follow an S-shape input-output transfer function with a trainable region-specific curvature. Activity decays with a trainable region-specific rate. The ordinary differential equation of the model is thus:

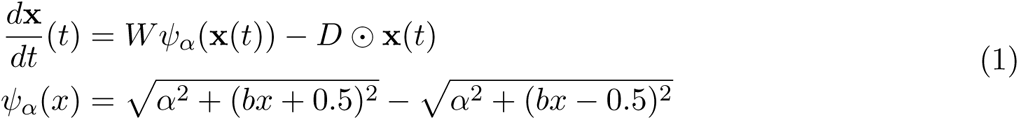

Here **x**(*t*) ∈ ℝ*^N^* is the neural activity hidden state at time *t*, where *N* is the number of parcels. *W* ∈ ℝ*^N×N^* is the connectivity matrix. The transfer function *ψ_α_* is applied element-wise with each region’s respective curvature parameter *α*. *D ∈* ℝ*^N^* is the decay and ⊙ indicates element-wise multiplication. *b* is another parameter controlling the shape of the transfer function, currently fixed as 20*/*3. To prevent overfitting and improve interpretability, we required the connectivity matrix *W* to be the sum of a sparse matrix *W_S_* and a low-rank matrix 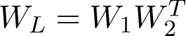, where *W*_1_*, W*_2_ ∈ ℝ*^N×k^* and *k < N* . Here we chose *N* = 200 (*k* = 72) as it achieves a balance between granularity and computational efficiency. However, we also repeated the analysis with *N* = 100 and *N* = 400 and obtained very similar attractor motifs.

We obtained **x**(*t*) from the preprocessed rfMRI data for each participant and session, with two runs combined. The ground-truth derivative Δ**x**(*t*) was computed using forward differentiation. Since HCP data utilizes an exceptionally fast TR, we performed two-point moving average smoothing on the derivative to reduce noise, i.e., Δ**x**(*t*) := [**x**(*t* + 2) *−* **x**(*t*)]*/*2. Removing the smoothing led to very similar results. We optimized *W*, *α* and *D* to minimize the difference between predicted derivative 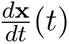 and ground-truth derivative Δ**x**(*t*) while enforcing sparsity of the connectivity using L1 regularization (Donoho, 2006). The loss function is defined as:

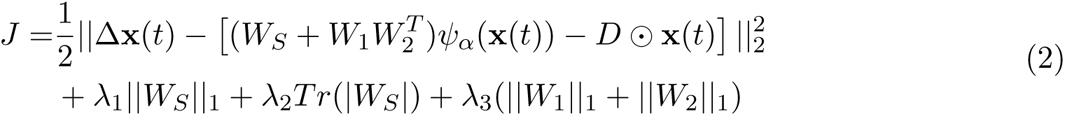

where *λ*_1_ = 0.075, *λ*_2_ = 0.2, *λ*_3_ = 0.05 are the regularization hyperparameters, *Tr*(*|W_S_|*) is the absolute sum of the diagonal elements of *W_S_*. Optimization was performed using gradient descent with Nesterov Accelerated Adaptive Moment Estimation (NADAM, (Dozat, 2016)). Parameters were updated after each minibatch of 300 random samples. We stopped the training at 5000 minibatches when the test-retest reliability of the parameters started to drop. To prevent the weights from being unnecessarily small due to regularization penalty, we performed an additional global rescaling of the parameters by fitting Δ**x** = *p_W_ Wψ_α_*(**x**) *− p_D_D* ⊙ **x**(*t*) with two scalar parameters *p_W_, p_D_* ∈ ℝ, and factored them into *W* and *D*. It’s worth noting that a 200-parcel MINDy model can be fit on a standard laptop within 15 seconds, enabling the analysis of the whole HCP dataset within a reasonable amount of time.

### 2.3 Model simulation and numerical analysis of attractors

For each model, we randomly sampled 120 initial conditions from the standard normal distribution. We also tried to sample from the data **x**(*t*), but the identified attractors were the same as long as the sample size is large enough. The dynamics (equation 1) was integrated for 1600 TRs using Euler’s method with step size equals to one TR. For some models with limit cycles that contained extremely slow ghost attractors, we prolonged the simulation until the state recurred after a full cycle. The stable equilibria were identified as follows: First, we defined a trajectory (simulation) as already converged to a stable equilibrium if *|x*(*t* + 1) *− x*(*t*)*| <* 10*^−^*^6^ for every parcel and every time point *T −*10 *≤ t ≤ T −*1, where *T* is the simulation time. The terminal state *x*(*T*) from all converged trajectories were than clustered together based on a simple Euclidean distance threshold of 0.1. The cluster centroids were extracted as the stable equilibria. Similarly, we defined a trajectory as already converged to a stable limit cycle if it approached and then left a small neighbourhood of the terminal state *{x | ||x − x*(*T*)*||*_2_ *<* 0.5*}* at least once. The interval during the last recurrence and the end of the simulation was considered as the period of the limit cycle, and the samples during this period was extracted to represent the limit cycle. We confirmed the validity of the method by visually inspecting the trajectories and identified attractors after dimensionality reduction using Principal Component Analysis (PCA), as depicted in Figure 1 and 2.

**Figure 1:**
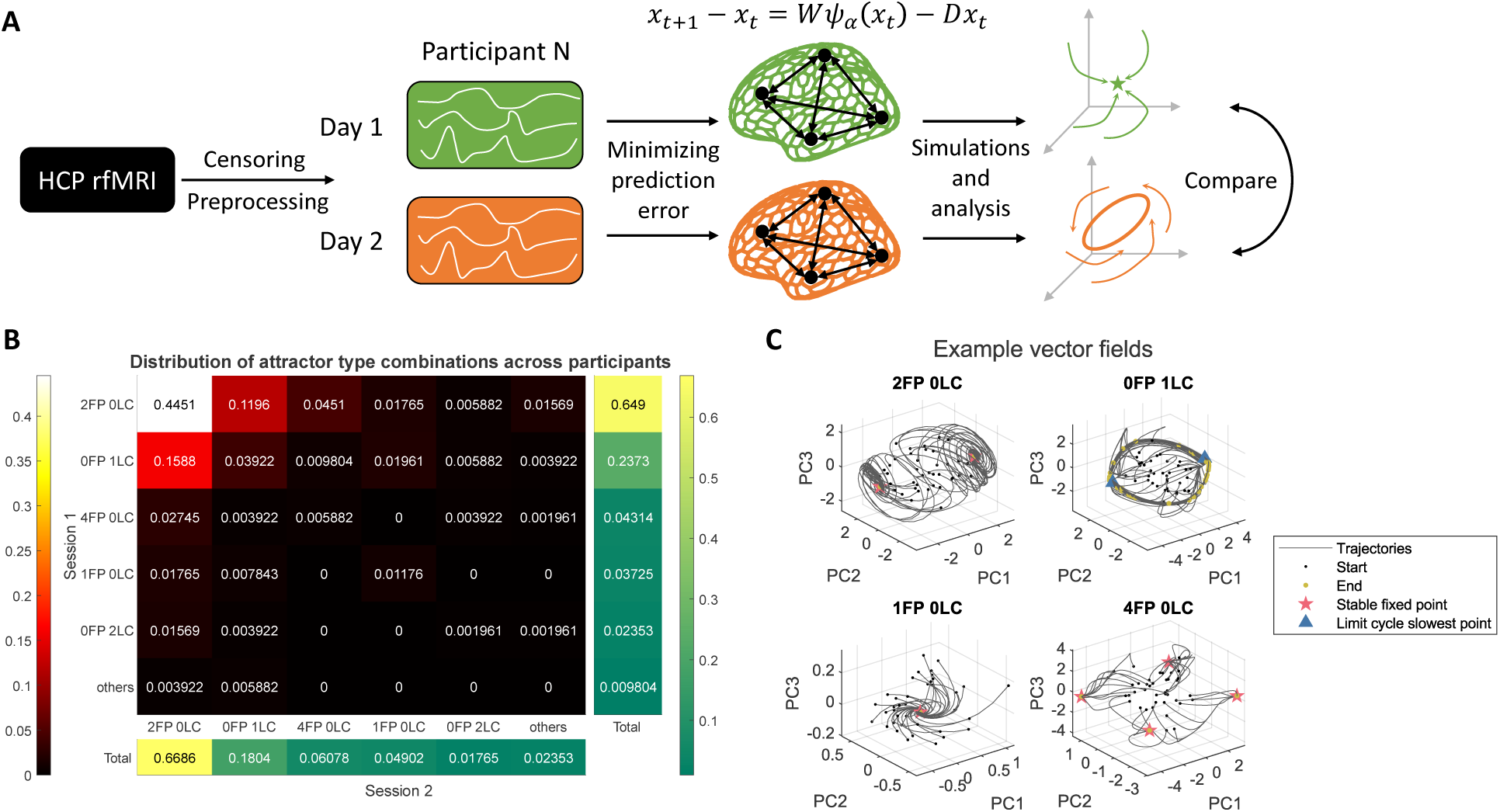
Nonlinear dynamical landscapes underlying individual rfMRI data. **(A)** Diagram of the analysis pipeline. **(B)** Distribution of dynamical landscapes across the participants. Numbers indicate the proportion of participants showing certain type of dynamics in session 1 (column index) and session 2 (row index). The types were defined using the number and types of attractors. ‘FP’ indicates stable equilibrium (fixed point) and ‘LC’ indicates stable limit cycle. Types with frequency less than 0.5% were grouped into ‘others’ (see Supplementary Figure 9). Marginal distribution for session 1 and session 2 are shown in the panel on the right and at the bottom respectively. **(C)** Simulated trajectories and obtained attractors from example models, projected onto the first three principal components (PCs) of the trajectories. Black and yellow dots mark the initial and final state of each trajectory. Numerically identified stable equilibria are denoted by red stars. The slowest points on the limit cycles were denoted by blue triangles (see Methods). Note that the distribution of final states (yellow dots) on the limit cycle reflects the relative size of the speed. The denser the distribution, the slower the dynamics. We used 120 simulations per model to identify the attractors but only showed 40 here for visualization.

**Figure 2:**
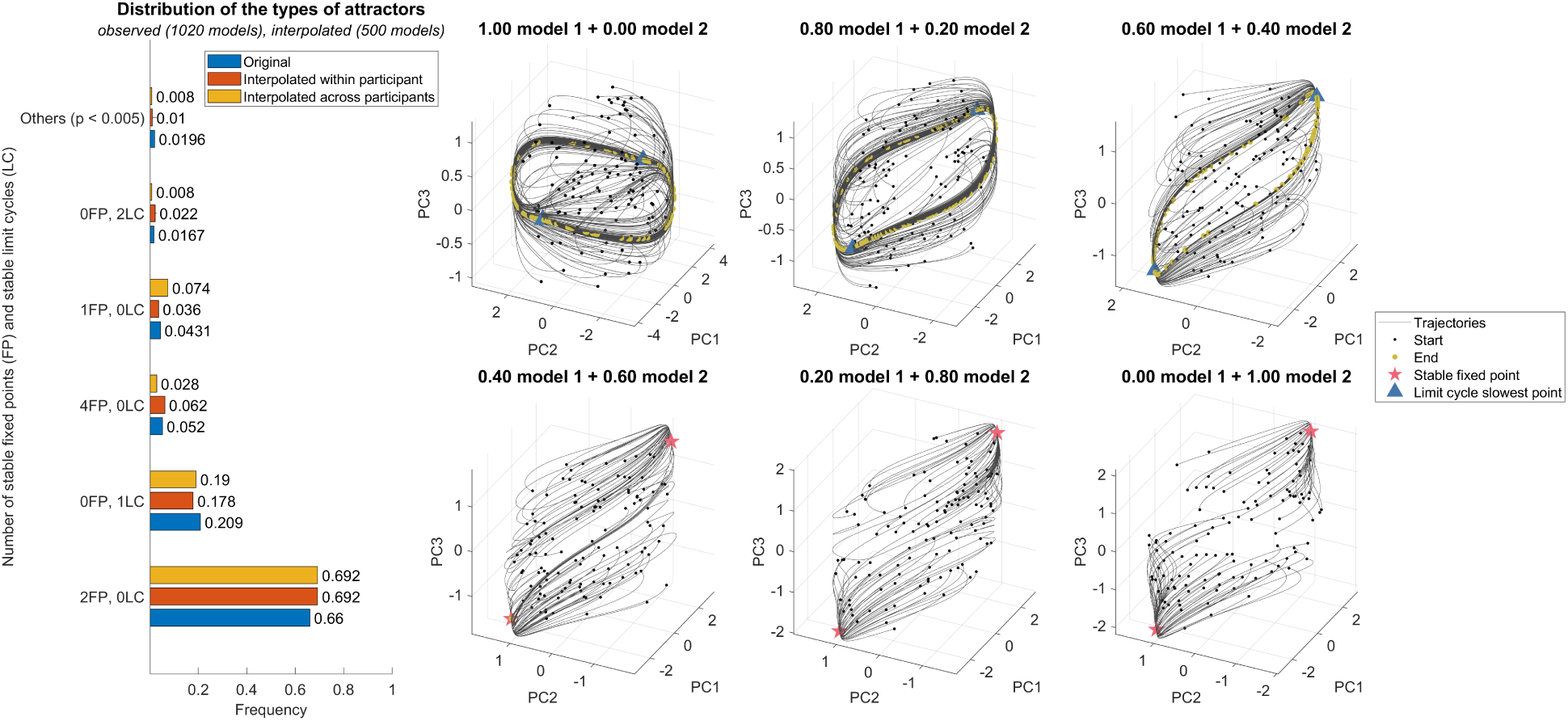
Observed taxonomy of dynamics were consistent with bifurcation-induced continuous spectrum. Left: Distribution of the type of dynamics in linearly interpolated models and empirically fitted models. The interpolated models were obtained by convex combinations of the dynamics from two models from the same participant in different sessions (‘within-participant’) or randomly sampled from all models (‘across-participant’). Right: Bifurcation process between two models of a same HCP participant. Trajectories and attractors were visualized in the same way as Figure 1. Note that the distribution of speed on the limit cycle (indicated by the distribution of final states) became more and more non-uniform as the first model bifurcated towards the second model.

We observed that the distribution of speed on the limit cycles was very non-uniform, sometimes varying by orders of magnitude. Therefore, we selected the slowest point on each limit cycle arg min**_x_**_(_*_t_*_)_ *||***x**(*t* + 1) *−* **x**(*t*)*||*_2_ as the ‘ghost’ attractors. Besides, for models with one limit cycle, due to the symmetry of the dynamics *f* (*−***x**) = *−f* (**x**), we added a pair of symmetric slowest points rather than a single one.

### 2.4 Bifurcation analysis

We induced a bifurcation between two models by creating convex combinations of the dynamics. Denoting the two model’s dynamics as **x**(*t*+1)*−***x**(*t*) = *f*_1_(**x**) and **x**(*t*+1)*−***x**(*t*) = *f*_2_(**x**) respectively, we construct a new model as **x**(*t* + 1) *−* **x**(*t*) = *αf*_1_(**x**) + (1 *− α*)*f*_2_(**x**), where 0 *≤ α ≤* 1 is the bifurcation parameter. We considered two ways to combine the observed models. In the within-individual case, *f*_1_ and *f*_2_ were the two models from the two sessions of a same participant. In the across-individuals cases, *f*_1_ and *f*_2_ were sampled from all models. We randomly sampled *α* from the uniform distribution in [0, 1] as well as a pair of models for 500 times, and extracted the attractors of these bifurcated models using the same numerical method described above. We then characterized the vector field by the number and types (equilibria or limit cycles) of attractors and compared their distribution across the bifurcated models and the fitted models in Figure 2.

### 2.5 Reliability analysis

We quantified the anatomical similarity of two attractors (and ghost attractors, same below) by the Pearson correlation between their coefficients (i.e., anatomical projections). As the number of attractors might differ across models, we defined the *dominant attractor similarity (DAS)* between two models as the maximum similarity between their attractors. The distribution of DAS between all pairs of models from different sessions was shown in Figure 3, separated by whether the two models come from a same participant and whether they have same type of dynamics (i.e., same number of stable equilibria and limit cycles).

**Figure 3:**
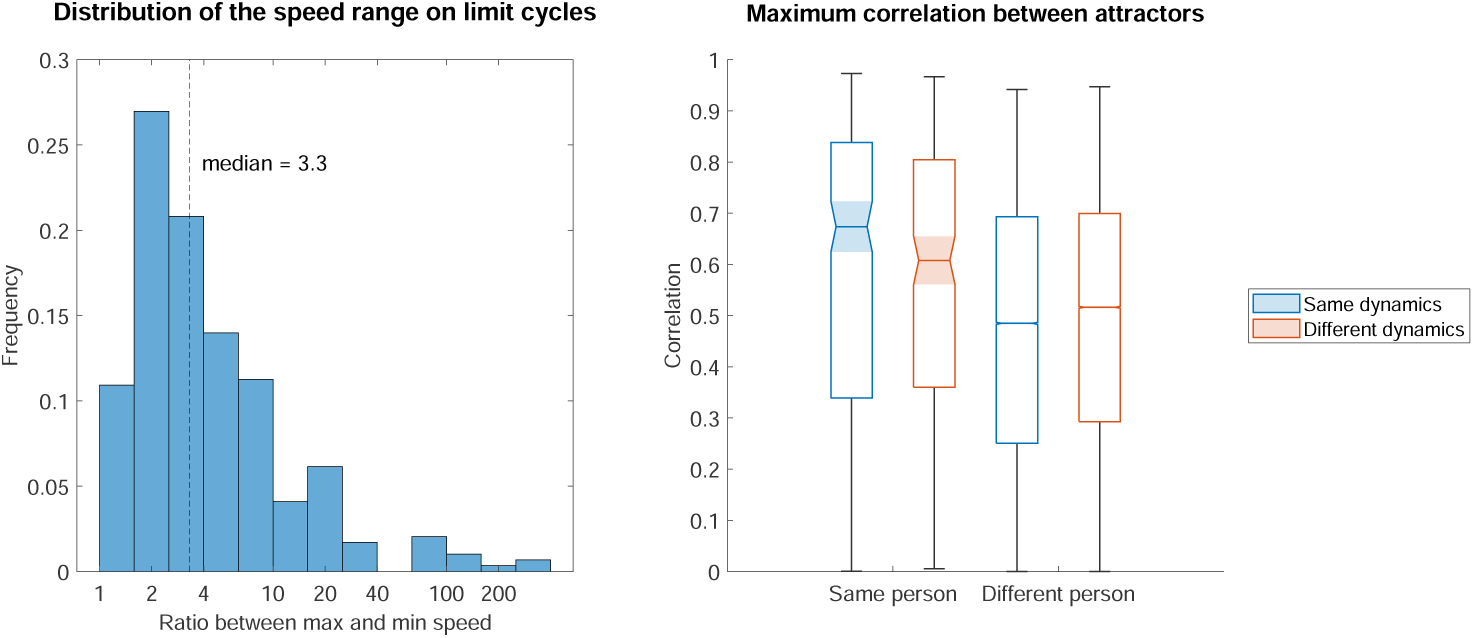
Attractors were more similar within than across individual regardless of changes in dynamical landscapes. Left: Distribution of the speed variability on limit cycles. For each limit cycle in all fitted models, the ratio between the maximum and minimum speed on the limit cycles was calculated. Right: box plot for the distribution of maximum attractor correlation between all pairs of models, separated by whether the two models come from a same person and whether they have the same type of dynamics (see Methods). Lines on the boxes indicate the maximum, first quartile, median, last quartile, and minimum of each distribution. Tapered, shaded region around the median indicates 95% confidence interval of the median.

### 2.6 Clustering analysis

We used *K*-means algorithm to cluster the anatomical location of all attractors and ghost attractors across all participants and sessions. We used cosine distance *d*(*x, y*) = 1 *−* cos ∠(*x, y*) for clustering and scaled the samples to unit norm for visualization. Using Euclidean distance produced almost identical clustering results. The number of clusters *K* is determined by the cluster instability index (Lange et al., 2004) with a candidate list of *K* ranging from 2 to 10. For each *K*, the data was randomly divided into two subsets for 30 times. We ran *K*-means algorithm on the first partition to obtained the cluster centroids, and used these centroids to classify the samples in the second partition. The classification result was compared to the results of directly running *K*-means on the second partition. A misclassification cost was computed after matching the labels using the Hungarian algorithm. This cost is then averaged across the 30 partitions and normalized by the null cost computed in the same way but with random labels, resulting in an instability index for each *K*. The local minimum of instability was selected as the number of clusters for the final clustering. We also repeated our analysis using either only the stable equilibria or only the ghost attractors on the limit cycles (Supplementary Figure 13, 14), and in each case both the clustering instability index and cluster centroids were very similar to the results in the main text.

## 3 Results

### 3.1 Model parameters captured reliable individual differences

We obtained 1020 MINDy models, one for each of two rfMRI scanning session associated with 510 HCP participants (Figure 1A). The models explained about 40% of variance of the derivatives in the test set (Supplementary Figure 5). The consistency of each parameter set (connectivity, curvature, decay) within individuals and across sessions was around 0.7-0.8. This quantity dropped to around 0.5-0.7 between individual, indicating that the obtained models were reliable and individualized (Supplementary Figure 7).

To perform an initial validity check that our models captured meaningful individual differences in the dynamics, we attempted to assess whether our obtained parameters can be connected to individual variation in cognitive measures. We performed Canonical Correlation Analysis (CCA) between the connectivity matrix of the models and the phenotypic measures available in the HCP dataset (Goyal et al., 2020; Smith et al., 2015). Interestingly, we obtained very similar results to (Smith et al., 2015), who used a connectivity matrix obtained via independent components analysis. We identified a single ‘positive-negative mode’ that was significantly correlated between the MINDy connectivity matrix and behavioral measures, and explained a significant proportion of variance for both. Post-hoc correlation found that this mode is most positively related to fluid intelligence, and most negatively related to substance use (Supplementary Figure 8). Therefore, the obtained MINDy models indeed capture reliable individual behavioral differences.

### 3.2 rfMRI embeds diverse nonlinear dynamics with nontrivial attractors

We next analyzed the asymptotic behavior of the neural trajectories. For this, we randomly initialized and forward simulated the models to obtain their limit sets and, specifically, reveal any asymptotically stable fixed points. As initially suggested in (Singh, Braver, et al., 2020), the dynamics captured in our obtained MINDy models were highly nonlinear, with nontrivial attractors and limit cycles found in the large majority of cases (Figure 1C). Less than 5% of models exhibited a single stable equilibrium at the origin (Figure 1B). It is worth noting that the observed non-trivial dynamics can not be simply attributed to any bias of our method, because MINDy models will actually recover trivial dynamics when fit on the noisy simulations of a stable linear system (Supplementary Figure 6). Most importantly, MINDy also correctly produced a globally attractive origin, rather than nontrivial attractors, when fit on noise with covariance and mean spectral power density that matches real data (Supplementary Figure 6). On the contrary, it is known that standard tvFC methods cannot disambiguate such noise and actual data (Laumann et al., 2017). Our findings thus support an interpretation of time-varying rfMRI activity as being most appropriately described emanating from a nonlinear dynamical system.

### 3.3 Induced bifurcations explained the heterogeneity of observed dynamics

Interestingly, although the obtained MINDy parameters were very reliable within each individual across sessions (Supplementary Figure 7), the number or type (fixed point or limit cycle) of attractors were still different in about half of all participants (Figure 1B). This finding indicates the existence of bifurcations, in which a small change in the model parameters results in a topological discontinuity of the dynamical landscape. Therefore, we hypothesize that resting state dynamics can be better described by a spectrum of possible dynamics that are sampled at each session, rather than a single monolithic brain state. To test the hypothesis, we constructed a parameterized ‘interpolated’ model as a convex combination of obtained models (i.e., (*γ*)Model_1_+(1 *− γ*)Model_2_), either within each participant (between the two models) or across all participants. If the hypothesis is correct, by varying *γ*, it should be possible to induce bifurcations. We analyzed the dynamics of the interpolated models and compared the distribution of the number and types of attractors with those of the original fitted models (Figure 2). The taxonomy of dynamics was very similar for fitted models and interpolated models, consistent with the idea that the dynamics we obtained are a reflection of whole brain dynamics that are undergoing bifurcations between topologically distinct vector fields.

### 3.4 Anatomically reliable individualized attractors marked signatures of bifurcations

Having established that the observed dynamics can be understood as bifurcating across a continuous spectrum, it is natural to ask how the brain can maintain stable or consistent function if the underlying dynamics are constantly changing. We thus hypothesized that there must be some anatomical commonality between the different obtained dynamics, leading to consistency in whole-brain activation patterns.

To probe this question, we looked for aspects of the dynamics that were relatively invariant to the bifurcations we observed. Most notably, we found that when the dynamics differed across sessions for a same person, they most commonly switched between having two equilibria or having one limit cycle (Figure 1B). This is consistent with the well-known infinite-period bifurcation (Keener, 1981). An infinite-period bifurcation begins with a limit cycle that contains a ghost attractor. As the bifurcation parameter changes, the ghost attractor becomes ‘infinitely slow’ (i.e., neural activity lingers near it for long periods of time) until eventually the bifurcation occurs and a stable node (along with an saddle node) gets created out of it (see Supplementary Figure 10 for an analytic toy model). In our data, when limit cycles were observed, the distribution of speed along them was highly non-uniform, sometimes varying by orders of magnitude (Figure 3, left). In our interpolated models, this form of bifurcation indeed occurs (Figure 2, right panel). Therefore, we hypothesize that the ghost attractors and point attractors provide a set of stable ‘operating points’ for the changing dynamics during rest.

If the session-to-session variability in fitted dynamics can be explained by such a bifurcation, we would expect that ghost attractors should be close to the stable equilibria in the other session, i.e., they should represent anatomically similar activation patterns. We thus defined ghost attractors as the slowest point on each limit cycle, and calculated the anatomical similarity between all (true and ghost) attractors. Because the number of attractors might differ across sessions, we defined the *dominant attractor similarity (DAS)* to be the maximum correlation over all pairs of attractors from the two models under consideration (see Methods). The DAS was higher within subject versus across subject (Figure 3, right), even when restricting the former to models showing different types of dynamics (e.g., two equilibria or one limit cycle) and the latter to models showing same type of dynamics (the second and the third boxes in the middle in Figure 3, right). Therefore, our results support the hypothesis that the resting brain is bifurcating between different dynamics, while maintaining a set of reliable attractors as operating points.

### 3.5 Attractors aligned with canonical resting state networks across population

Next, we examined whether these attractors could be interpreted from a functional standpoint. Interestingly, the DAS was high (around 0.5) even between different participants (Figure 3), indicating the existence of consistent patterns across the whole population. Therefore, we clustered the locations of the attractors across all participants and sessions, where the number of clusters *K* were selected according to cluster instability index (Lange et al., 2004) (see Methods, Supplementary Figure 11). We obtained near perfect cluster stability only for *K* = 4, apart from the less interesting solution *K* = 2. Note that the attractors always exist in pairs (see Methods) so *K* = 2 represents only one pattern and its opposite (Supplementary Figure 12).

The individual attractors within each cluster aligned very well with the cluster centers (Figure 4). Even more interestingly, the activation patterns were highly modular, respecting the functional network organization defined in the (Schaefer et al., 2018) atlas, even though the model fitting process was completely agnostic to parcel labeling. In particular, one large cluster was dominated by the activation of DMN and FPN. Another cluster was dominated by the FPN and dorsal/ventral attention networks. The other two clusters showed the opposite activation profiles. The clusters and network organization in combination accounted for more than 40% of the total variation across all participants, sessions, and parcels (Supplementary Figure 15). The activation pattern was better explained by functional network organization rather than the spatial proximity of parcels on the cortical surface (Supplementary Figure 16). Furthermore, the attractor clusters emerged regardless of whether we only included the equilibria, the limit cycle ghost attractors, or both (Supplementary Figure 13, 14), supporting the hypothesis that they represent the reliable activation dynamics of resting state networks.

**Figure 4:**
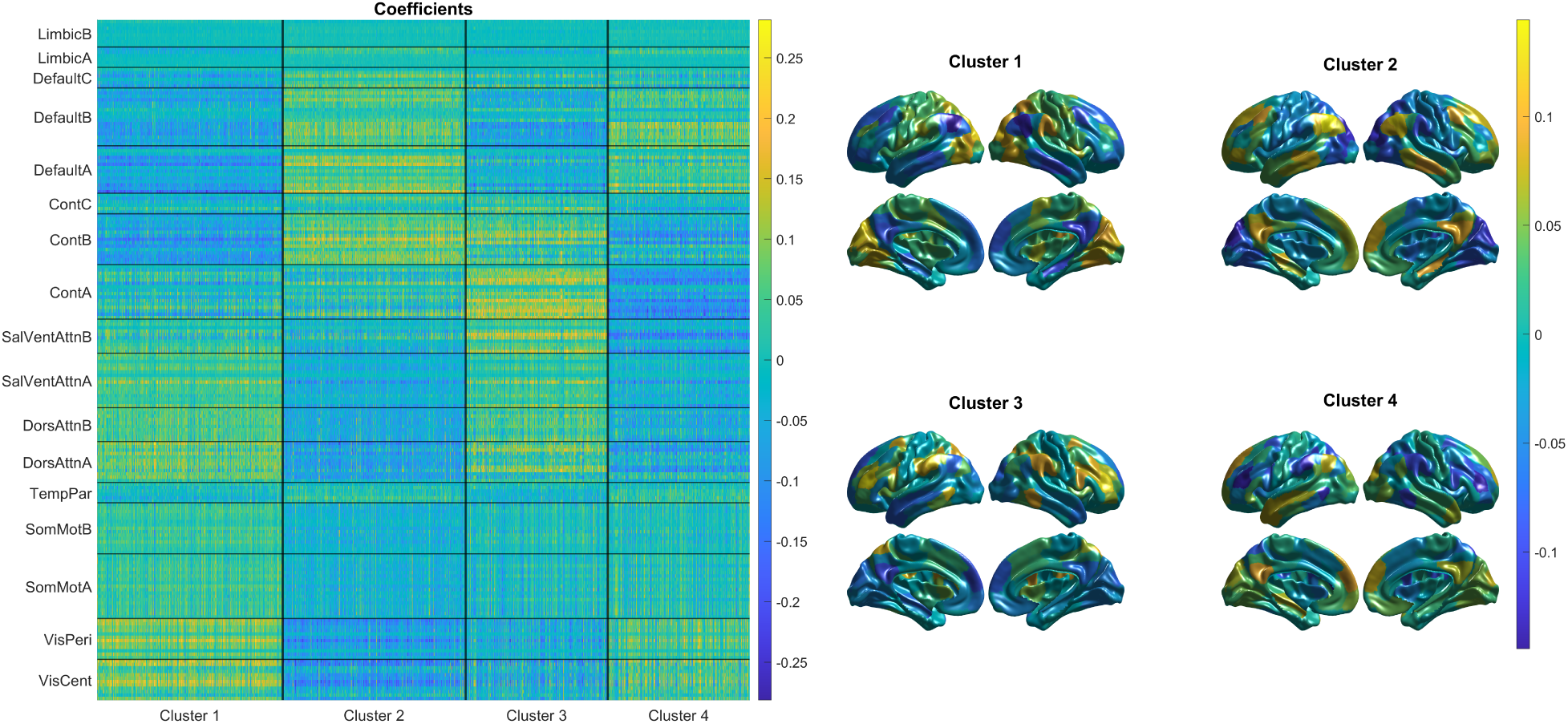
Attractors aligned with canonical resting state networks across the population. Left: locations (coefficients) of all attractors. Each row corresponds to one parcel and each column corresponds to one attractor. Rows are sorted according to the (Schaefer et al., 2018) atlas and columns are sorted according to cluster assignments. Thick horizontal and vertical lines separates the functional networks and clusters, respectively. Attractors are scaled to unit norm for visualization purpose. Right: cluster centroids visualized as the activation patterns over the cortex.

## 4 Discussion

In this study, we adopted a nonlinear dynamical systems modeling framework to analyze how resting state brain dynamics varied across individuals and time. We found that resting state brain activity embedded diverse nonlinear dynamics that included nontrivial attractors rather than a single globally stable equilibrium at the mean. Interestingly, the dynamics reliably varied between individuals. Moreover, instead of being stationary, the dynamics underwent bifurcations across different scanning sessions even within the same individual. Furthermore, the observed spectrum of dynamics could be fully recovered through induced bifurcations between fitted models. Consistent with such bifurcations, the attractors and ghost attractors were anatomically reliable within and between individuals. These attractors were organized into distinct clusters that reflected the activation of different functional brain networks, particularly the DMN, the FPN, and the dorsal/ventral attention networks. Using the formal language of dynamical systems and bifurcation theory, our results shed light on the connection between nontrivial fluctuations in resting state activity, the organization of functional brain networks, and stable individual differences. We provide a modeling and analysis framework that describes the individualized nontrivial dynamics in rfMRI, while maintaining interpretability and tractability. A variety of dynamical features can be derived from the models for future brain-phenotype association studies. Our results also enable model-based interventions into brain dynamics, which might hold great potential for individualized treatments of brain disorders and even cognitive enhancement (Singh et al., 2022).

### 4.1 Nontrivial attractors and resting state networks

Resting state brain activity is traditionally treated as a unimodal stationary process fluctuating around a stable mean. However, recent studies have begun to explore nontrivial recurring patterns in resting state dynamics (Abreu et al., 2020; Liu & Duyn, 2013; Vohryzek et al., 2020). Mostly notably, time-varying functional connectivity (tvFC, also called ‘dynamic’ FC) studies have suggested that the resting brain is traversing multiple states associated with distinct brain-wide association patterns (Lurie et al., 2020). However, it is debated whether these recurring patterns indeed represent nontrivial fluctuations, or merely capture the snapshots of the trivial fluctuations around the mean (Laumann et al., 2017). Due to these challenges, it has been questioned whether the new insights we gain from these techniques are relatively marginal compared to the elevated difficulties in analysis and interpretation (Nozari et al., 2023).

We argue that the controversies over non-trivial fluctuations reflect the lack of mechanistic interpretations that can be derived from short-time-windowed methods typically utilized to estimate tvFC. Dynamic-systems-based modeling provides a more generative explanation to the observed dynamics and statistics. Phase portraits of the fitted models reveal insights about the resting brain dynamics without sacrificing the ease and interpretability of analysis. Here, by using a nonlinear, individualized, and fully trainable dynamical systems modeling framework, we provide evidence for nontrivial fluctuations in rfMRI dynamics. Most importantly, when fitted on stationary noise with FC and a mean power spectrum that match real data, our model correctly recreated a monostable dynamic system, while tvFC has been demonstrated to produce spurious nontrivial states in this situation (Laumann et al., 2017). Therefore, the nontrivial attractors consistently observed in our fitted models are less likely to be explained by a bias inherent in the method. Compared to other modeling studies, our model relies on fewer assumptions about connectivity structure. Instead of assuming that white matter density is a good surrogate of functional coupling at the mesoscale level, which has been challenged by some studies (Barttfeld et al., 2015), we posit a ‘sparse plus low rank’ structure with all the weights directly optimized towards empirical fMRI timeseries. Very interestingly, even without prior constraint on the functional network structure, the attractors that emerged from the models still exhibited highly modular activation patterns that respect the organization of functional networks. Such convergence of evidence is further supported by the fact that, the MINDy effective connectivity between localized parcels and the functional connectivity between distributed ICA-defined networks, despite using very different node types and connectivity measures, encoded highly similar information about cognitive individual differences as revealed by CCA (Supplementary Figure 8). Furthermore, unlike most previous work that has relied upon population-level models, we were able to show that the nontrivial attractors are not only consistent across the population, but also test-retest reliable within each individual.

It has long been hypothesized that nontrivial attractors represent ‘functional’ states that can be spontaneously traversed during rest, as if exploring the repertoire of operating points (Deco, Jirsa, & McIntosh, 2013). Although there has been some theoretical work focused on the functional relevance of resting state attractors (Kurikawa & Kaneko, 2013), empirical evidence has been scarce. Our work supports this hypothesis by showing that nontrivial resting state attractors reflect selective activation of functional brain networks, and contain reliable individual differences. It will thus be very interesting to extend the MINDy framework to task states, in order to analyze how such nontrivial attractors might be engaged in cognitive computation. Similar to DCM methods (Friston et al., 2019), we can couple the MINDy recurrent dynamics with an input term representing the task control signal. The interaction between task demand and inherent dynamics and attractors can thus be potentially characterized through control-theory-based analysis. Such analysis will shed light on how the resting brain ‘prepares’ stable motifs for cognitive computation, advancing of our understanding of the mechanistic link between the resting state and cognition.

### 4.2 Bifurcations and the critical brain

We presented three lines of evidence that resting state brain dynamics are not only nontrivial, but also bifurcating. First, the topology of the dynamic landscapes changed across sessions even when the controlling parameters remained highly reliable, consistent with the definition of bifurcations. Second, we induced bifurcations between fitted models and recovered the full spectrum of the dynamics observed, confirming that the fitted models can be understood as samples from such a continuous spectrum encompassing several bifurcations. Third, we identified anatomically reliable attractors or ghost attractors on a cycle, consistent with the prediction of an infinite-period bifurcation. Our results thus provide a richer description of both the invariants and changes in resting state dynamics, showing that even though the statistical outputs (FC) of two datasets might seem similar, they could be supported by distinct but still intimately related dynamics.

The most interesting conclusion from our analysis is that the resting state brain is highly sensitive to modulation, in which a small perturbation can bifurcate it towards various different dynamics. To illustrate this point, we first showed that even though the correlation between the parameters of any two models was high (around 0.6), we still observed at least eight different kinds of dynamics, in the sense of different numbers or types of attractors. Moreover, we found that although the parameter correlation within each participant was even higher (around 0.75), the dynamics were still different between sessions for almost half of all participants.

We suggest that the sensitivity of resting brain dynamics might provide one mechanism for cognitive flexibility, as the brain can easily bifurcate from the resting state towards various different dynamics that might be advantageous for different cognitive computations. If that is the case, the task state brain dynamics should be more rigid than resting state, and should vary across tasks according to the computation required. In line with this idea, it was found that task-driven input reduced the variability of whole-brain dynamics compared to rest (Ito et al., 2020; Ponce-Alvarez et al., 2015). Further, the global connectivity of the frontoparietal network was found to systematically vary across 64 tasks, with more similar connectivity for tasks that require more similar computation (Cole et al., 2013). Most interestingly, it has been found that the similarity between rsFC and task-specific FC positively correlated with task performance across multiple tasks, and the similarity between rsFC and task-general FC can be related to fluid intelligence (Schultz & Cole, 2016), indicating that cognitive ability is related to how efficiently resting state dynamics can transform into a variety of different task dynamics. Our study provides a more mechanistic framework that can capture the changes in the spectrum of generative dynamics, not just the statistics of such dynamics (i.e., FC). Extending the MINDy framework to task contexts will thus provide a strong test for such hypothesis.

Our results also relate to the idea of criticality in the brain. Note that the word ‘criticality’ has been used in at least two different ways in neuroscience. In the broad sense, criticality refers to the emergence of slow, large-amplitude (scale-free) fluctuations in a dynamical system that is close to losing its stability, without specifying the kind of instability to which it is transformed (Cocchi et al., 2017). In a narrow sense, such transformation is restricted to that between an ordered state (stability) and an unordered state (chaos) (O’Byrne & Jerbi, 2022). Our results support criticality of rfMRI dynamics in the broad sense rather than the narrower sense. We found that our models show either a stable low-activity state with no nontrivial attractors, or an unstable low-activity state with nontrivial attractors, suggesting the existence of supercritical bifurcations where a high-activity attractor emerged ‘above’ the low-activity attractor as the latter loses its stability. Such bifurcations have been shown to give rise to slow, scale-free fluctuations (Cocchi et al., 2017). In previous models of whole-brain dynamics, it has been found that the criticality associated with such bifurcations improved the response sensitivity to external stimuli (Deco, Jirsa, & McIntosh, 2013). However, critical dynamics are not always optimal for all tasks. For example, the sensitivity to inputs also reduces the reliability of the response (Fagerholm et al., 2015). Therefore, it is proposed that instead of staying critical, brain dynamics should reverberate between multiple regimes near criticality (Wilting et al., 2018). Our results support this hypothesis with novel evidence that the brain traverses near criticality, potentially balancing the computational advantages of each regime across different computations.

If the resting brain resides in this critical regime that can be easily modulated into different dynamics, one interesting question is how the brain might implement such modulation to utilize different dynamics. On a longer timescale, such as the dynamics we obtained here across a 30-minute resting state scan, neuromodulator systems might be the best candidate. It is well established that neuromodulators can affect brain connectivity and whole-brain dynamics (Shafiei et al., 2019; Shine et al., 2018). The arousal system might play a particularly important role in the fluctuations of resting state dynamics (Laumann et al., 2017). On a shorter timescale, such as during the execution of cognitive tasks, goal-directed top-down modulation might also bifurcate the dynamics. Theoretically, such bifurcations might relate to the proactive control mode in the dual mechanism framework for cognitive control (Braver, 2012). In contrast to reactive control, proactive control refers to the active maintenance of goal-related information and biasing cognitive computations even before cognitively demanding events occur. In parallel, goal-directed bifurcation might transform the dynamical landscape to bias the neural processing even before receiving cognitive inputs. Therefore, combining MINDy modeling with cognitive tasks as well as neuromodulatory manipulations holds great potential to further our understanding about the dynamical flexibility of the brain.

### 4.3 Limitations and future directions

In this study, we characterized the variable nature of resting state dynamics by comparing across participants and scanning sessions. However, such variation inevitably interweaves with measurement and modeling error. A stronger test for the non-stationarity of resting state dynamics requires comparing the dynamics across different periods within a single scanning run. Such analysis is difficult to applied to the HCP which only has 15-minute long scans, roughly as much as the data needed to obtain a reliable MINDy model (Singh, Braver, et al., 2020). However, it can be done on datasets with as long as 30 contiguous minutes of rfMRI data, such as the Midnight Scanning Club (Gordon et al., 2017). Another approach is to use EEG or MEG data which have much lower dimension and much higher sampling rate. It might be even possible to update the model parameters in real time for EEG/MEG data (Singh et al., 2023).

There are also limitations associated with the MINDy model used in the analysis. The models did not include an intercept/bias term, and the dynamics are anti-symmetric with respect to the neural hidden states (see Methods). Such assumption was made according to the excitation-inhibition-balance principle (van Vreeswijk & Sompolinsky, 1996) and has been adopted in resting state DCM studies too (Razi et al., 2017). In such models, the zero vector (corresponding to the mean of the data) is always an equilibrium (though not necessarily stable) and nontrivial attractors (if any) must exist in pairs. Extending MINDy to task contexts by introducing a task-related input can break such symmetry. Another limitation of the current method is the ability to account for the variations in haemodynamics. Here, we estimated the hidden neural states by a noise-aware deconvolution of BOLD signal with the canonical haemodynamic response function (HRF) (Friston et al., 1998). However, it has been suggested that HRF varies significantly across brain regions and individuals (Aguirre et al., 1998). Although MINDy parameters have been demonstrated to be robust against HRF variations (Singh, Braver, et al., 2020), it is unclear how much the emergent dynamics will be influenced, given the observation that the dynamics were sensitive to parameters. Extending our analysis with region-specific HRF (Singh, Wang, et al., 2020) is a natural step to follow. Another direction is to use a biologically more detailed regional dynamics model (Friston et al., 2019). Currently, the recurrent dynamics within each brain region is modeled as a simple self-excitation (or inhibition) with an exponential decay. Despite being computationally more tractable, such a model might not capture the full dynamics within a region, especially the interactions between sub-populations (Singh et al., 2023). Multi-scale modeling and EEG-fMRI data fusion might be a possible way to improve biological specificity while maintaining computational efficiency.

From a cognitive neuroscience perspective, the current study demonstrates how the novel lens of dynamical systems and attractor landscapes can be utilized for theory and analysis regarding the relationship between intrinsic dynamics (i.e., present during resting states) and task-related cognitive computations. Future studies can try to associate individualized resting state dynamical motifs with cognitive traits and task performance using either traditional correlational analysis, or more interestingly, by adapting the MINDy framework to analyze how dynamics and attractors change between resting and task states. As a mechanistic alternative to FC and tvFC analysis, our framework can also generate new insights into the dynamical changes associated with different dimensions of cognitive variation, including states of consciousness (e.g., sleep, meditation, psychedelics), developmental stages, or dysfunction associated with psychiatric and neurological disorders.

## 5 Data and code availability

The rfMRI data is available at Human Connectome Project’s website. The MINDy modeling toolbox is available at this repository. The preprocessing scripts are available at this repository. All analysis scripts are available at this repository.

## 6 Author Contributions

**R.C.:** Conceptualization, Methodology, Software, Formal analysis, Data curation, Writing - original draft, Visualization. **M.S.:** Methodology, Software. **T.S.B.:** Conceptualization, Writing - review & editing, Supervision, Project administration, Funding acquisition. **S.C.:** Conceptualization, Writing - review & editing, Supervision, Project administration, Funding acquisition.

## 8. Acknowledgements

R.C. wants to thank Janine Bijsterbosch and Geoffrey Goodhill for helpful discussions. Portions of this work were funded by NIMH 1R21MH132240-01 and NINDS R01NS130693.

## 9 Supplementary materials

### 9.1 Validation of MINDy modeling

#### 9.1.1 Goodness of fit

We tested the prediction accuracy of the fitted models in three cases: the training data, the data from the same participant but in the other session, and the data from a random participant in the same session. The R squared was around 0.4 for all three cases, with the training accuracy being the highest, the within-person transfer being the second and across-person transfer being the third (Figure 5). Therefore, our models indeed captured the individuality of the dynamics.

**Figure 5:**
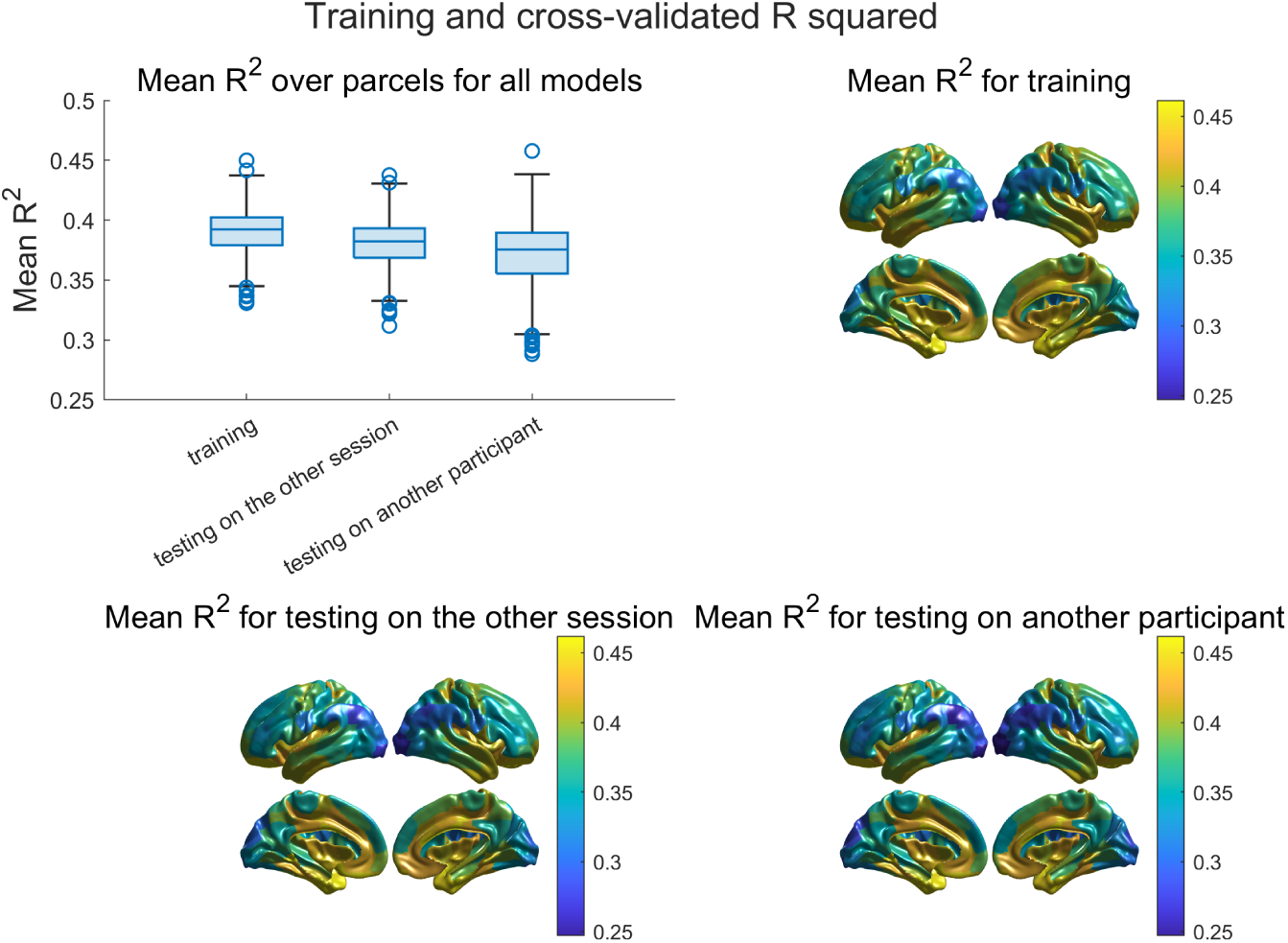
Goodness-of-fit and cross validation accuracy. Top-left: distribution of the mean R squared for all models when predicting the training data, the data from the same participant in the other session, or the data from another randomly selected participant. Top-right: Distribution of mean R squared over all models when testing on training data. Bottom-left and bottom-right: similar plots for testing on the other session within participant, or testing on another participant’s data.

#### 9.1.2 Surrogate data simulations

To make sure that the nontrivial dynamics observed in fitted models were not simply due to methodological bias, we investigated whether MINDy can correctly capture the trivial dynamics in closely matched surrogate datasets. We tried two different surrogate simulation schemes and fit MINDy on these simulations using the same hyperparameters as in the main text. The first method follows (Laumann et al., 2017), generating a dataset from noise (without any dynamics) but preserves the covariance (thus FC) and mean power spectrum of the rfMRI data. In (Laumann et al., 2017), it was found that the tvFC method generated indistinguishable results for real data and such surrogate data. However, our model correctly produced a monostable dynamic system without nontrivial fluctuations (Figure 6). The second method fit MINDy on the noisy simulations of a closely-matched linear system. This linear system, referred to as ‘linear MINDy’, replaces the nonlinear activation function of MINDy with its best linear approximation. The ‘linear MINDy’ model was fit to rfMRI data using the same hyperparameters and loss function as actual MINDy, thus capturing the statistics of the data and also maintaining the ‘sparse plus low-rank’ connectivity structure. In fact, we found that the connectivity and decay parameters of ‘linear MINDy’ is highly correlated with the actual MINDy model fitted on the same data. After fitting the linear model, we simulated the model with additive noise. The magnitude of the noise was set to the root of mean squared error during the fitting of the linear model. The noisy simulations thus represent a close match of the true dataset but generated from inherently linear (and monostable) dynamics. We then fit (nonlinear) MINDy models on these simulations, and the models correctly reproduced a monostable dynamic system with no nontrivial attractors (Figure 6).

**Figure 6:**
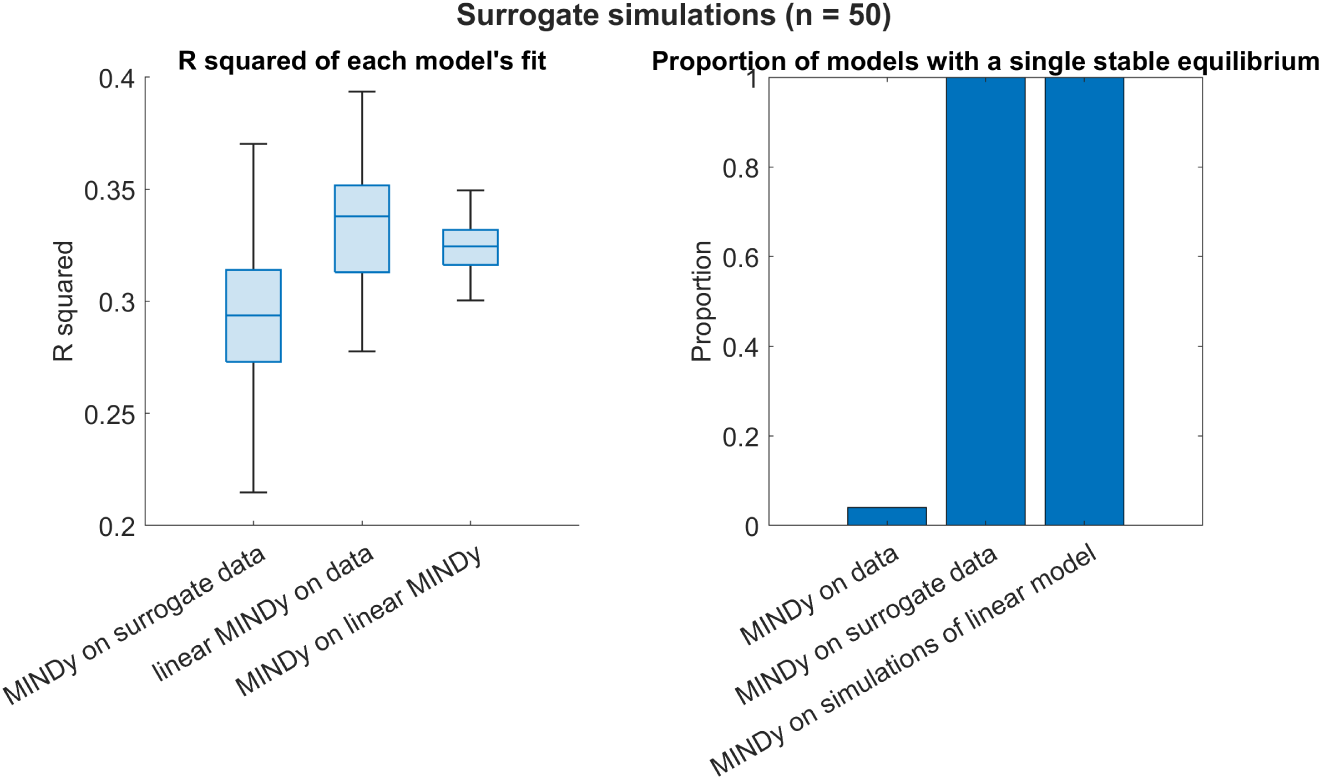
Fitting MINDy on surrogate data revealed trivial dynamics. Left: R squared for fitting MINDy on surrogate data; fitting ‘linear MINDy’ on rfMRI data; and fitting MINDy on the noisy simulations of ‘linear MINDy’ models. Right: the proportion of models showing a single stable equilibrium (at the origin), for MINDy models fitted on rfMRI data, on surrogate data, and on the noisy simulations of ‘linear MINDy’ models.

### 9.2 Reliability and behavioral correlation for MINDy parameters

#### 9.2.1 MINDy parameters were individualized and consistent across the population

We characterized the consistency of model parameters across all participants, as well as within each participant. We computed the correlation of each set of parameters (connectivity, curvature or decay) between each pair of models. For each participant, we quantified the similarity between their two models as well as the mean similarity between their models and all other models. Results indicate that the parameters were highly consistent across the population, and even more within each participant (Figure 7, top-left). Next, for each parameter (e.g., one entry in the connectivity matrix), we calculated its intraclass correlation coefficient (ICC), which is the correlation of its value between the two sessions across all participants. ICC characterized the reliability of the individual differences in each parameter. We observed a large set of connectivity parameters with high ICC (Figure 7, top-right; also note that the connectivity is sparse so a lot of entries have low ICC). The ICC for curvature and decay parameters were acceptable, around 0.5 (Figure 7, bottom).

**Figure 7:**
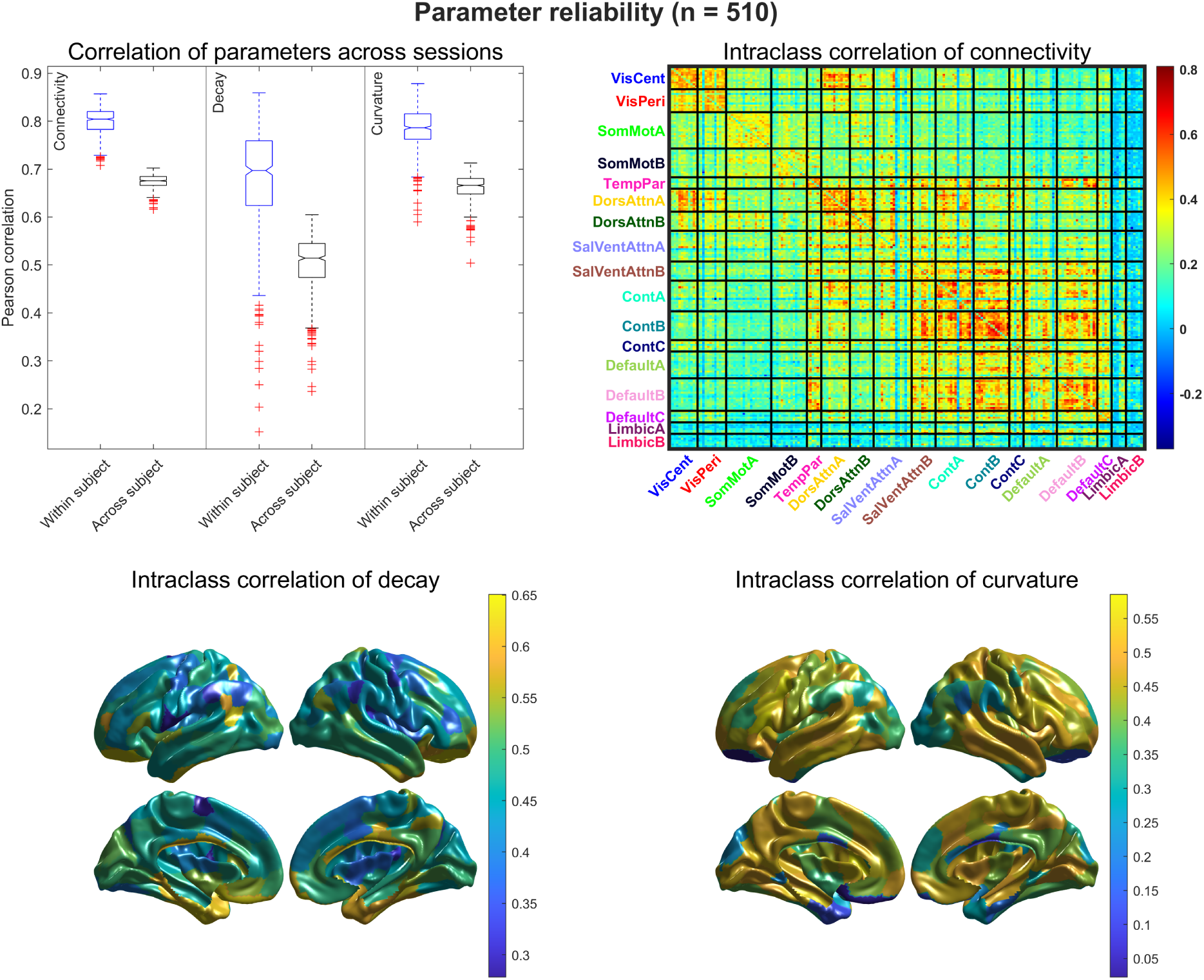
Obtained parameters were individualized and consistent across the population. Top-left: Correlation between sets of parameters (connectivity, curvature or decay) from models from different sessions. ‘Within subject’ is the correlation between the two models from a same participant, while ‘across subject’ is the mean similarity with all other models. Top-right: intraclass correlation coefficient (ICC) of each connectivity parameter. Text labels indicate the functional networks, separated by the black thick lines. Bottom: ICC for decay and curvature parameters, respectively.

#### 9.2.2 MINDy parameters encode cognitive differences

Here, we conducted a canonical correlation analysis (CCA) between the connectivity matrices of fitted models and the phenotypic measures in the HCP. The connectivity matrices from the two sessions were averaged within each participant before entering the analysis. We used the scripts provided by (Goyal et al., 2020) which extends (Smith et al., 2015) to the whole HCP dataset. The connectivity matrices and subject measures were first projected to their first 100 principal components to reduce dimensionality. Then, CCA was carried out to identify the directions to which the projection of the connectivity data and phenotypic data are maximally correlated across the population. We identified a unique pair of such directions (modes) with statistically significant correlation (1000 permutations, Figure 8, top right). Further, this connectivity mode and phenotypic mode explained a significant amount of variance in their data respectively (Figure 8, middle and bottom right). Post-hoc correlation between the phenotypic mode and all phenotypic measures revealed that this mode is more related to fluid intelligence and substance use (Figure 8, left). Interestingly, our results are very similar to the original finding in (Smith et al., 2015) even though we are using very different node types (parcels vs. ICA networks) and connectivity measures (effective vs. correlational).

**Figure 8:**
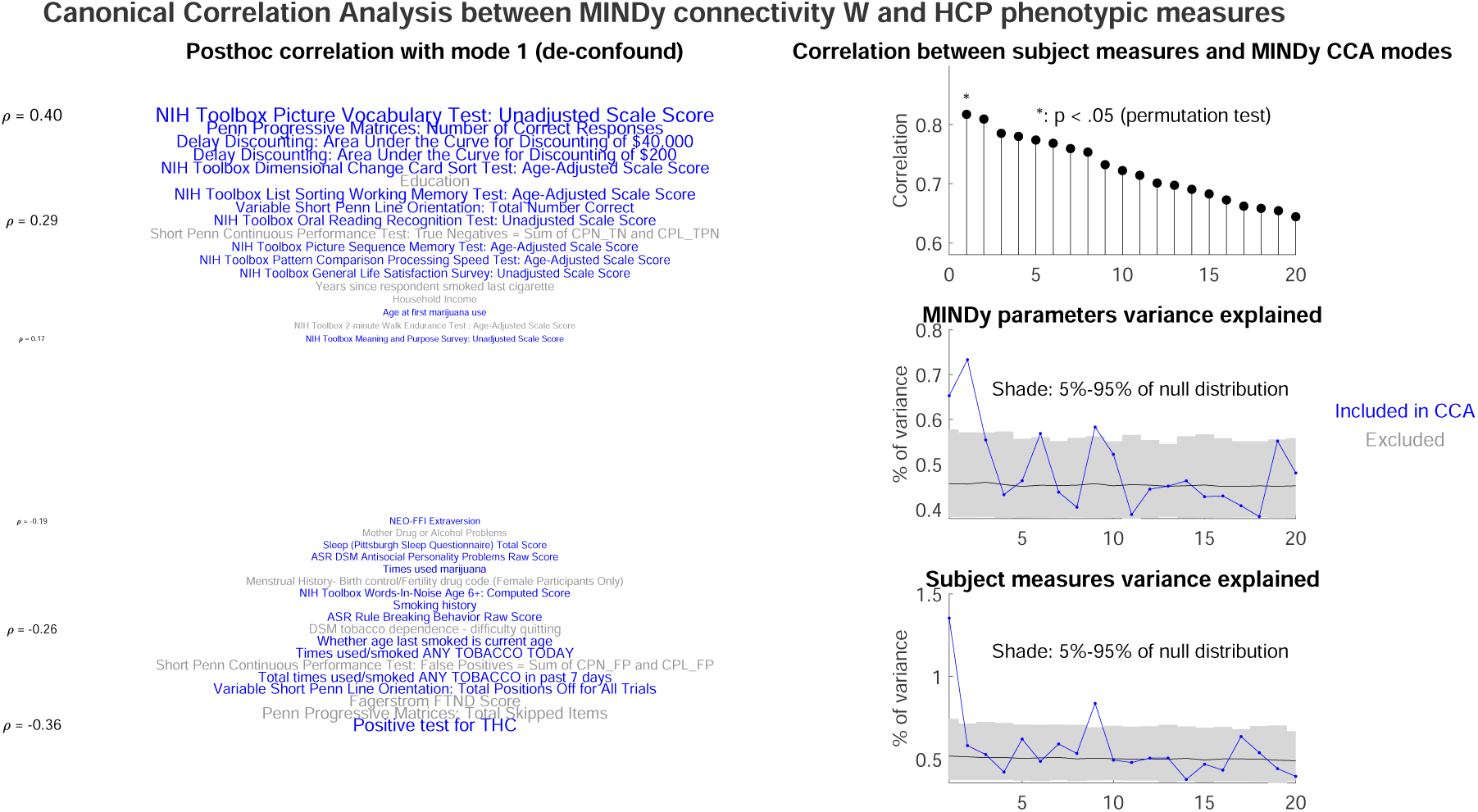
MINDy parameters encoded reliable individual differences. Please compare with Figure 1 in (Smith et al., 2015). Left: Post-hoc correlation between the behavioral mode identified by CCA and the phenotypic measures. We listed the most correlated measures with font size scaled by the correlation. Note that the Y axis is ordinal but not scalar. Top-right: correlation between CCA-identified connectivity and behavioral modes. Statistical significance is determined by permutation test with 1000 permutations (same for other panels). Mid-right: Variance of the connectivity explained by the CCA modes. Shaded region indicates the null distribution with 1000 permutations. Lower-right: variance of the behavioral measures explained by the CCA modes.

### 9.3 More examples of model dynamics

Here we show more examples of the dynamics observed in the fitted models (Figure 9). The three except the top-left one were grouped into ‘others’ type in the main text because of their rare occurrence.

**Figure 9:**
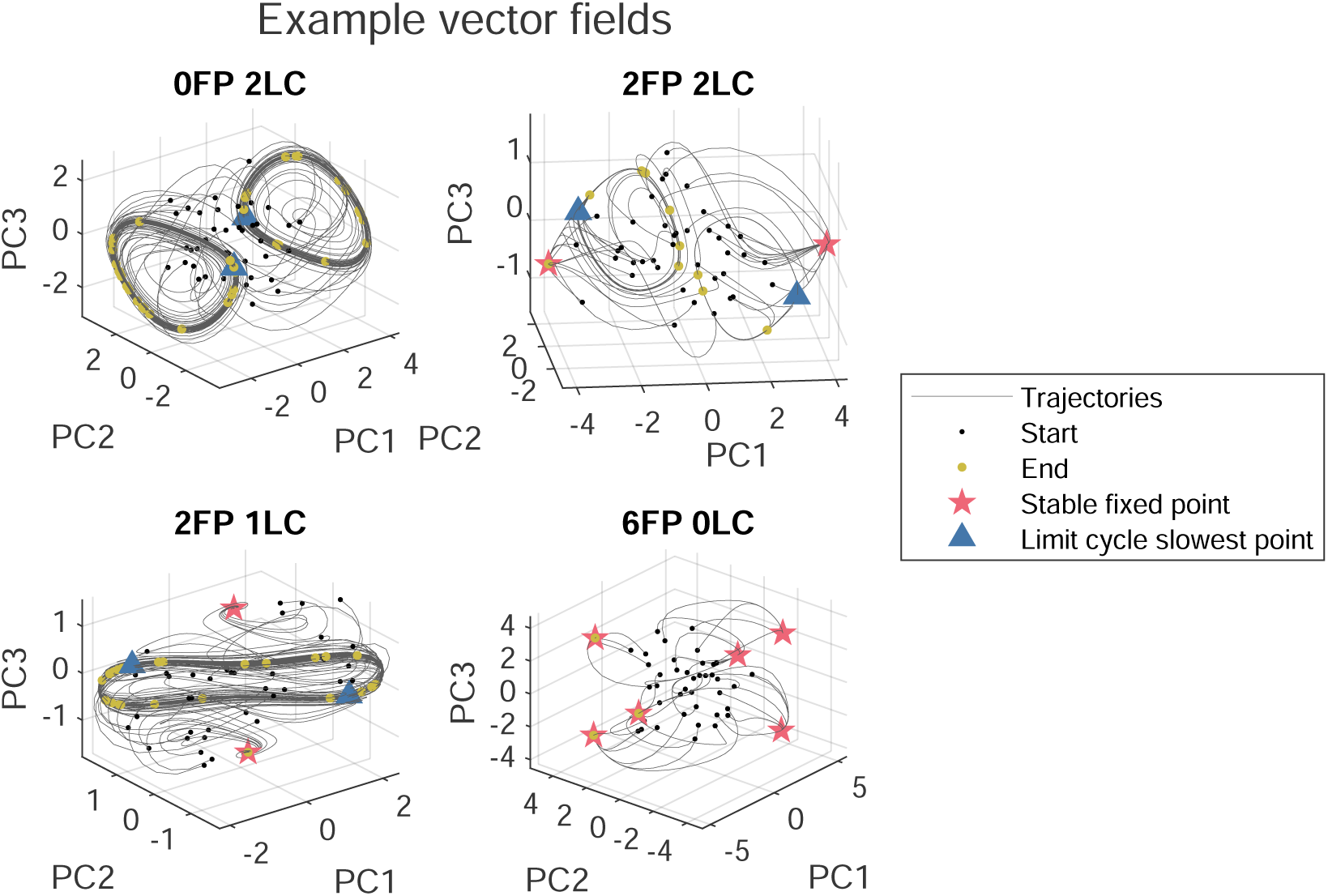
More examples of dynamical landscapes. See Figure 1 in main text.

### 9.4 Toy model for infinite-period bifurcation

Here we show a toy model for infinite period bifurcation mentioned in the main text. The model dynamics are written in polar coordinates (but shown in Cartesian coordinates in Figure 10) as:

**Figure 10:**
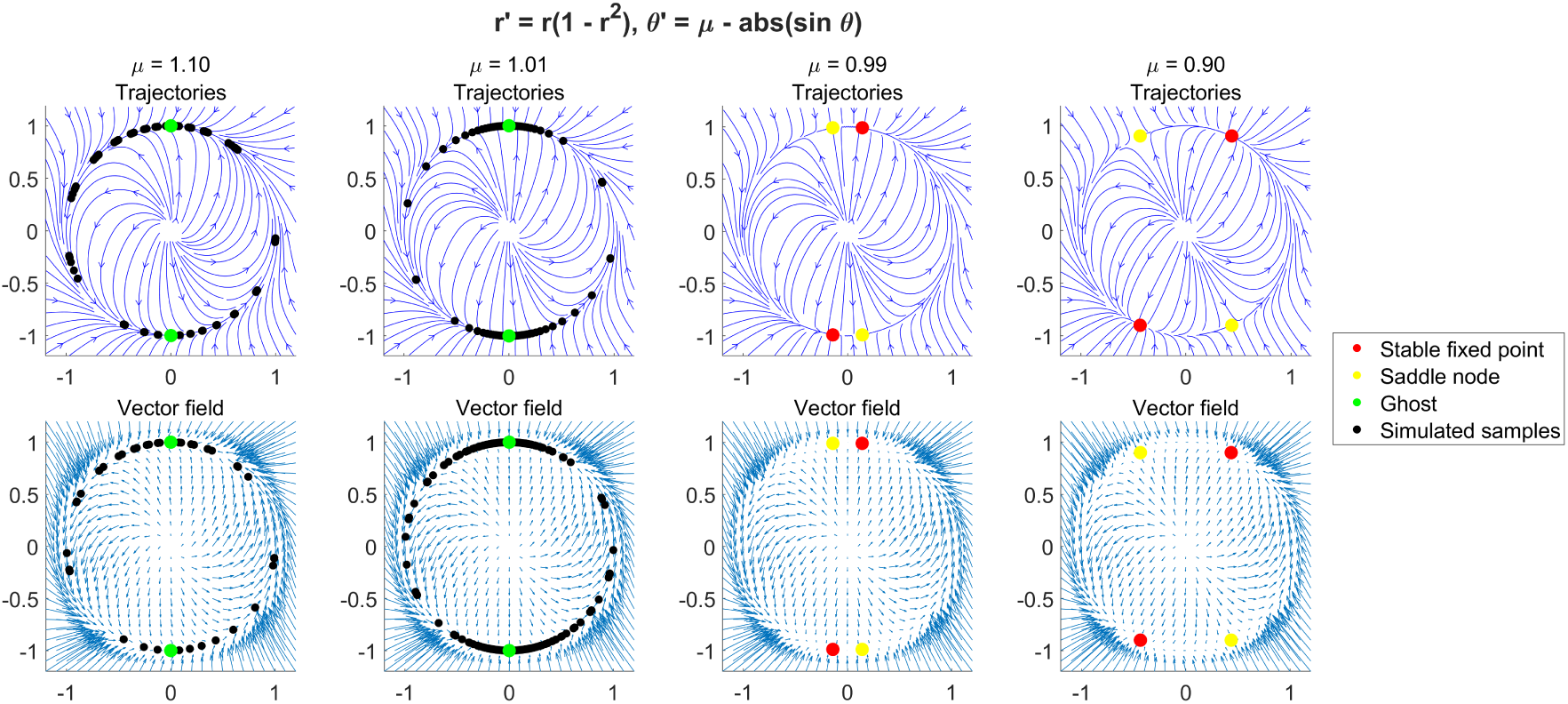
Toy model for infinite period bifurcation. Top row: trajectories of the models with different value for the bifurcation parameter *µ*. Red, green and gray dots indicate stable equilibria, saddles and ghost attractors respectively. Black dots are the simulated samples on the limit cycle. Bottom row: vector fields of the models.

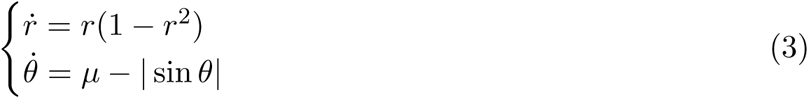

where *µ* is the bifurcation parameter. The infinite-period bifurcation happens when *µ* equals one. When *µ >* 1, the system shows a stable limit cycle and an unstable equilibrium at the origin. As *µ* approaches 1, the speed distribution on the limit cycle becomes more and more extreme and a ghost attractor emerges at (0, 1) on the limit cycle. When *µ* equals 1, the ghost attractor dissolves into a pair of equilibria, one stable and one unstable.

### 9.5 Clustering of attractor patterns

#### 9.5.1 Selection of the number of clusters

We used K-means algorithm to cluster the attractors and select the number of clusters *K* using the cluster instability index (see Methods for more details). We found that regardless of whether we use only the stable equilibria, only the ghost attractors, or both, we got near perfect clustering stability only for *K* equals to two or four (Figure 11).

**Figure 11:**
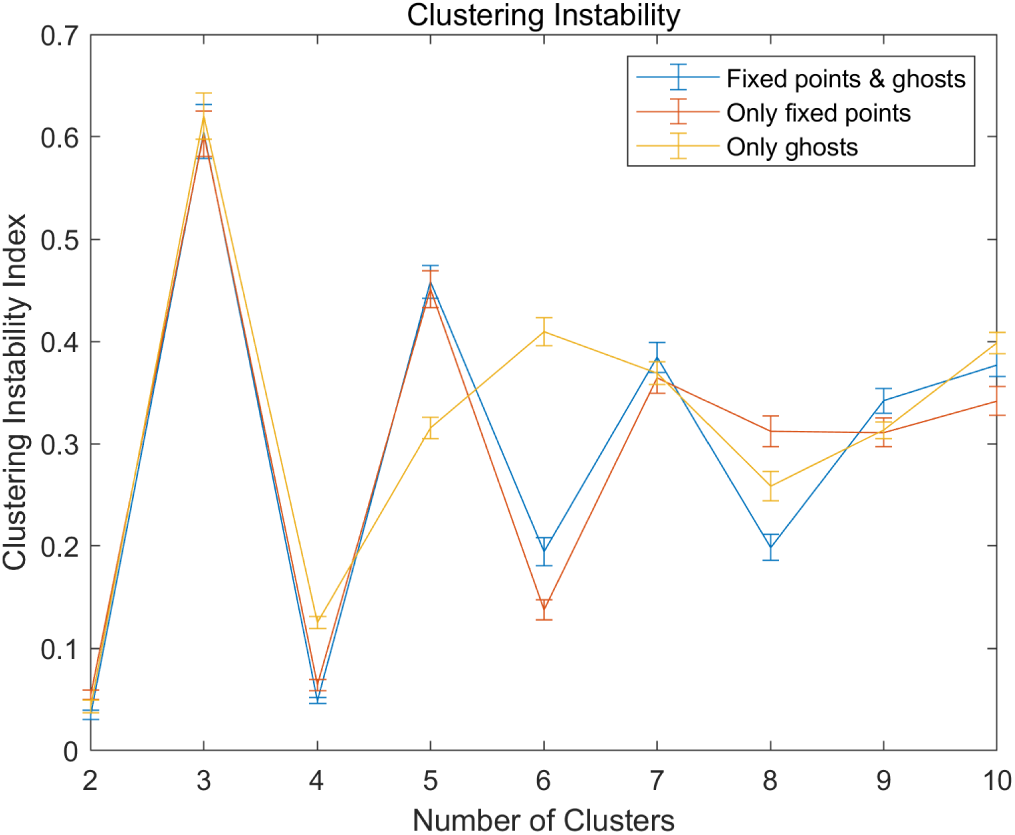
Stable clustering solution for two and four clusters.

#### 9.5.2 Two cluster solution

We show the two-cluster solution in Figure 12. In this case the two clusters are the reflections of each other. One of them showed strong activation for the DMN and FPN, while the other show strong activation for the visual and the dorsal/ventral attention networks.

**Figure 12:**
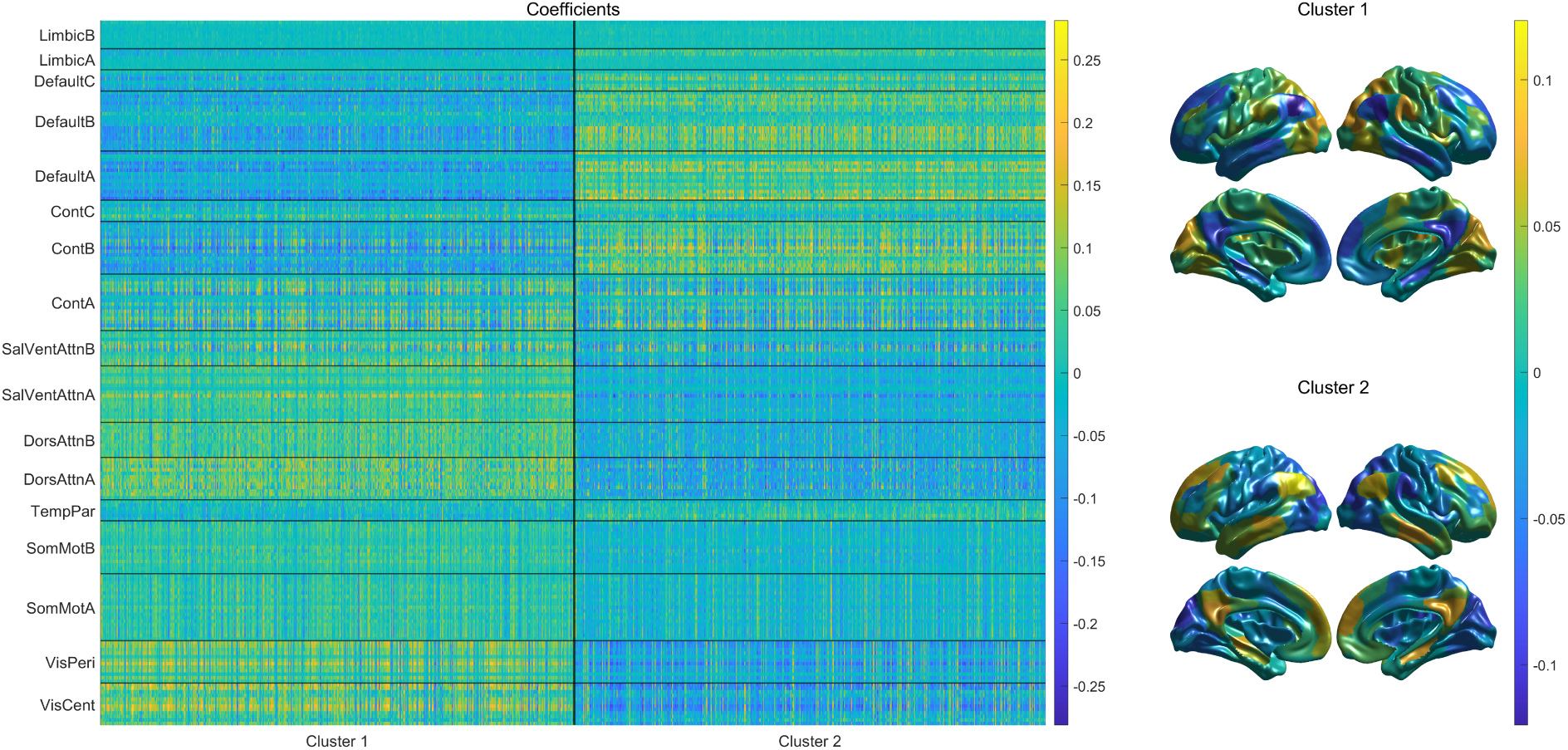
Two-cluster solution of the attractor clustering.

#### 9.5.3 Clustering results with only stable equilibria or only ghosts

In the main text we clustered all stable equilibria and ghost attractors together. Here we show the results using only stable equilibria (Figure 13) and only ghost attractors (Figure 14) respectively.

**Figure 13:**
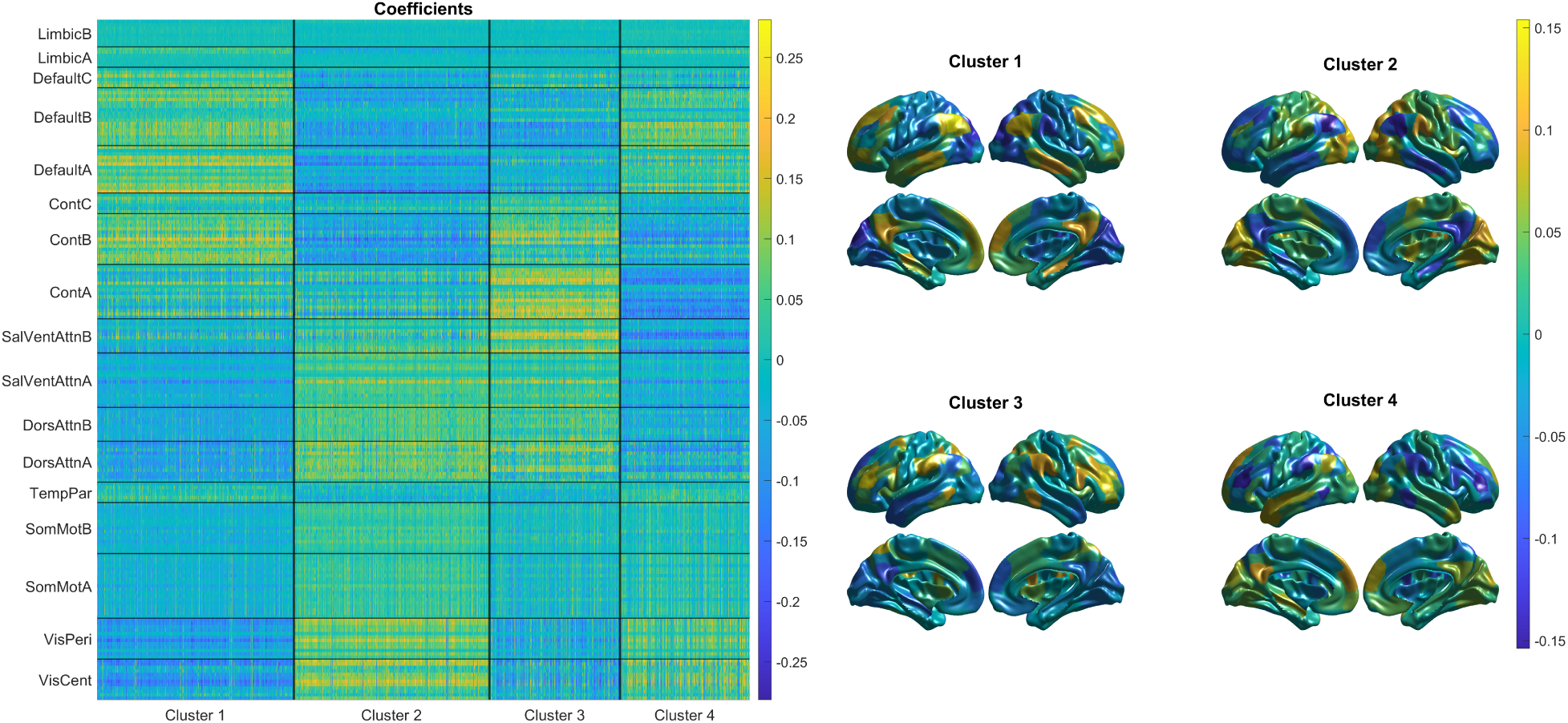
Attractor clustering with only the stable equilibria.

**Figure 14:**
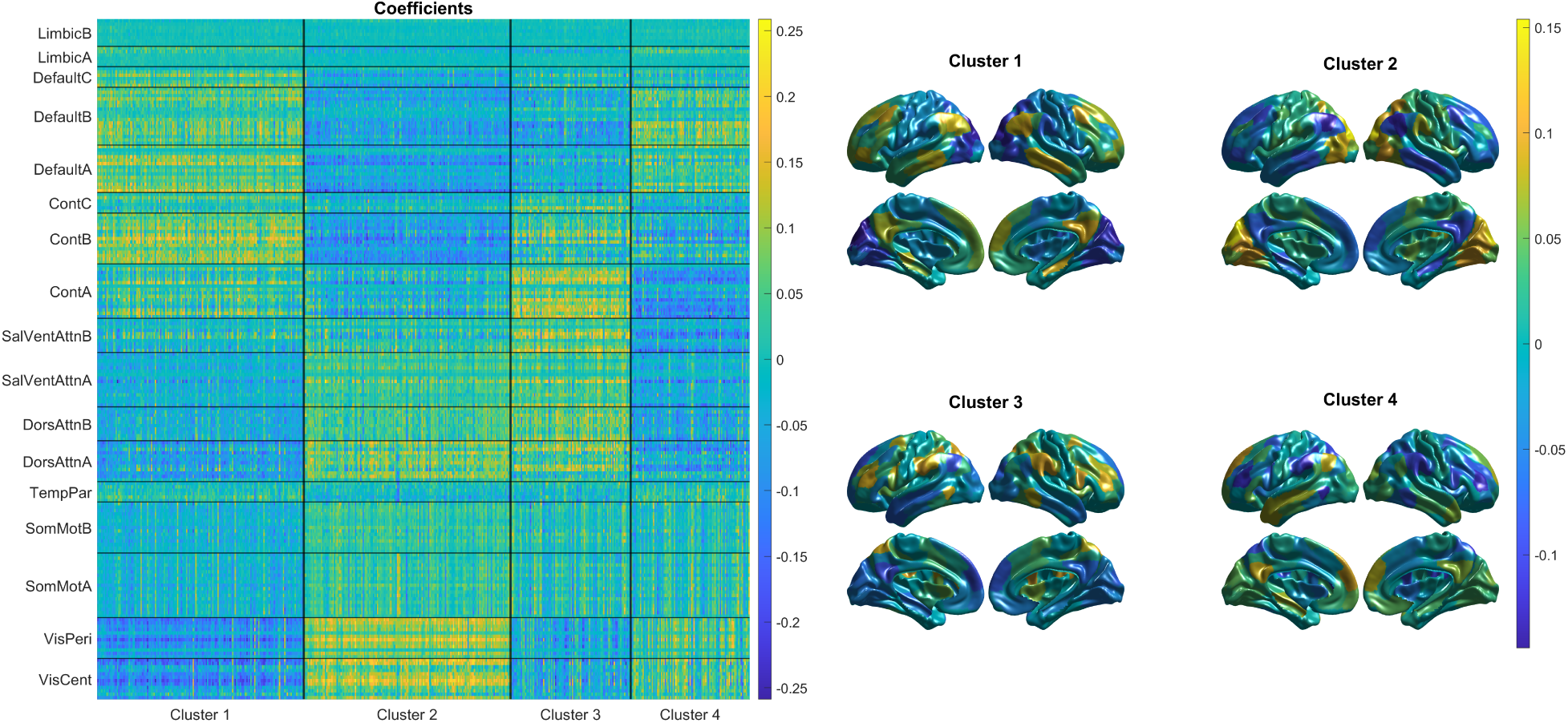
Attractor clustering with only the limit cycle slowest points.

## 10 Parcel activation in attractors

### 10.1 Parcel activation follows network and cluster structure

We analyzed the activation of parcels across all attractors with a mixed-effect model: *activation ∼ 1 + cluster + network + cluster:network + (cluster|network:parcel)*. We computed the hierarchical (type I) sum of squares explained by each term (Figure 15). The main effects of cluster and network were small while their interactions explained over 40% of total variation, indicating that (1) the same brain network showed very different activation in different clusters (or equivalently, that each cluster is associated with the strong activation of different sets of networks); and (2) parcel activation was mostly determined by the combination of attractor clusters and functional brain network structure. The random effect of parcels explained about 15% of variation, indicating that heterogeneity still exist across the parcels within each network. The error sum of squares was smaller than the total variation explained by the modeling, suggesting high consistency across participants and sessions.

**Figure 15:**
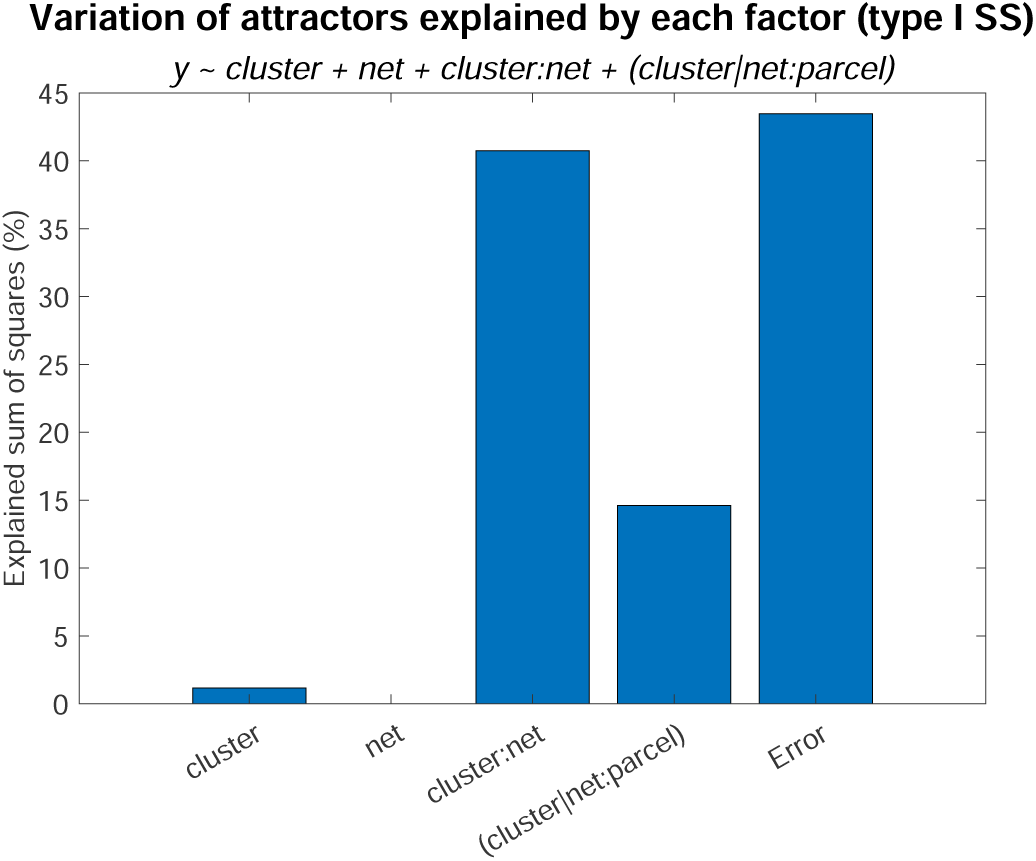
Variance of parcel activation explained by attractor clusters and functional networks.

### 10.2 Functional rather than spatial organization explains data better

To show that the activation was driven by functional network segmentation rather than the spatial proximity between parcels, we calculated the Pearson correlation between the activation patterns of each pair of parcels across all attractors. The correlation coefficient was Fisher-transformed into a Zr statistic (Figure 16, top-left). We modeled this similarity matrix by the combination of the spatial proximity (negative cortical distance) between parcels and their functional network assignment. The distance between all cortical vertices along the surface were calculated using the surface geometry file from HCP and MATLAB’s graph distance function, and then averaged within the two parcels under consideration. All inter-hemisphere entries were excluded since the distances were undefined. The functional network assignment similarity was set to one if two parcels belong to the same network as defined by the 17-network atlas in (Schaefer et al., 2018), and zero otherwise. We then predicted the activation similarity using negative cortical distance, functional network assignment and their interactions. A hierarchical sum of squares analysis showed that network organization explained 15% of total variation even after excluding the effect of spatial proximity (which explained must less variance, Figure 16, bottom-right, first column). Therefore, the attractors indeed reflected the organization of functional brain networks over and above the spatial configuration of cortical regions.

**Figure 16:**
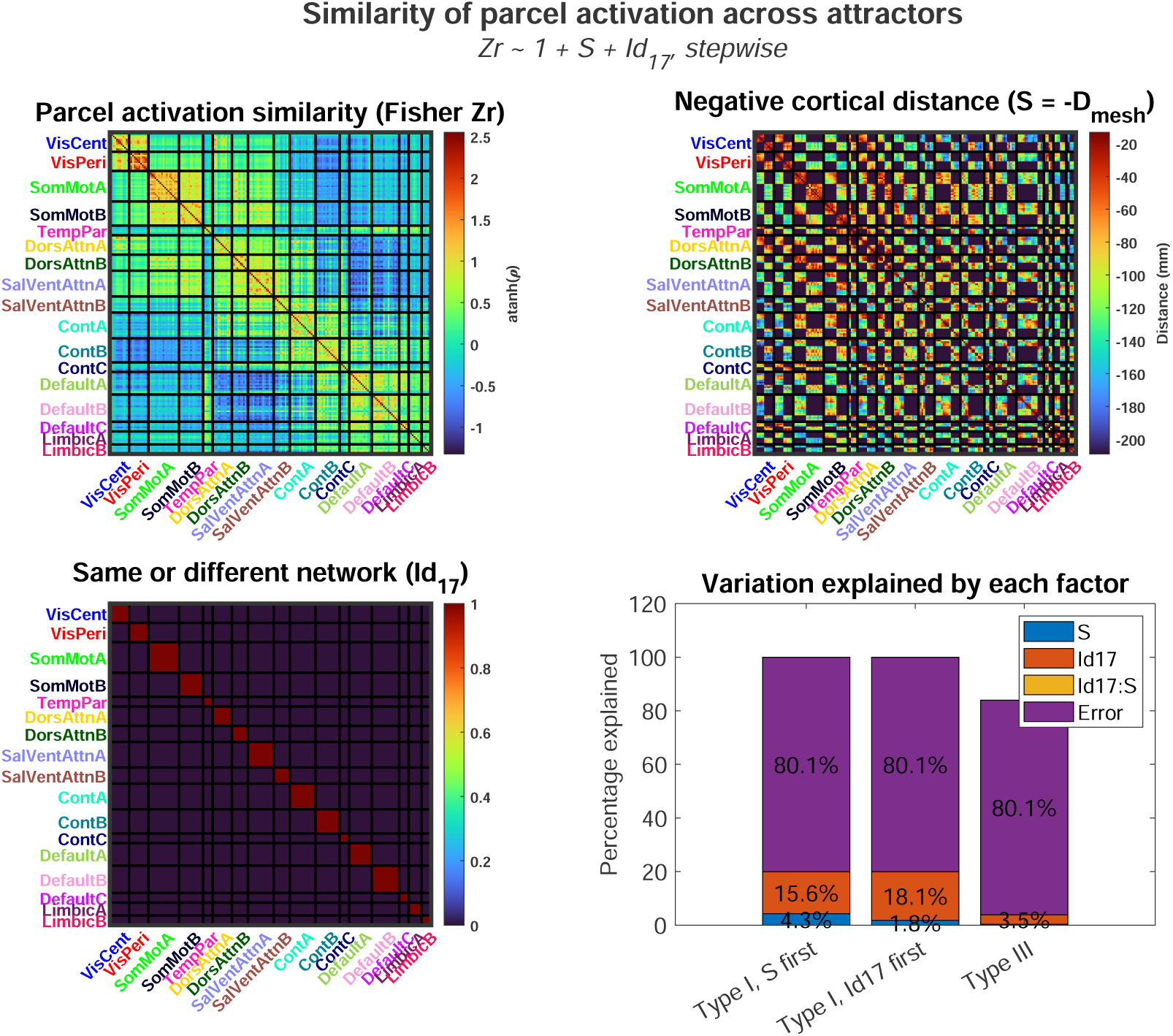
Parcel activation similarity across all attractors. Top-left: similarity between all parcels’ activation across all attractors. Similarity is quantified by Pearson correlation followed by Fisher’s transformation. Top-right: the negative of the distances between parcel centroids along the cortical surface. Inter-hemisphere entries were omitted. Bottom-left: similarity between parcels based on (Schaefer et al., 2018) functional network segmentation. Similarity is one if the two parcels belong to the same network and zero otherwise. Bottom-right: variance of parcel activation similarity explained by distance or network structure. Column one: type I (hierarchical) sum of squares (SS) where cortical distance precedes network structure (see the text). Column two: type I SS where network structure precedes cortical distance. Column three: type III SS. The SS for the interaction between distance and network structure is less than 1%.

